# Inverse sensitivity analysis of mathematical models avoiding the curse of dimensionality

**DOI:** 10.1101/432393

**Authors:** Ben Lambert, David J. Gavaghan, Simon Tavener

## Abstract

1

Biological systems have evolved a degree of robustness with respect to perturbations in their environment and this capability is essential for their survival. In applications ranging from therapeutics to conservation, it is important to understand not only the sensitivity of biological systems to changes in their environments, but which features of these systems are necessary to achieve a given outcome. Mathematical models are increasingly employed to understand these mechanisms. Sensitivity analyses of such mathematical models provide insight into the responsiveness of the system when experimental manipulation is difficult. One common approach is to seek the probability distribution of the outputs of the system corresponding to a known distribution of inputs. By contrast, inverse sensitivity analysis determines the probability distribution of model inputs which produces a known distribution of outputs. The computational complexity of the methods used to conduct inverse sensitivity analyses for deterministic systems has limited their application to models with relatively few parameters. Here we describe a novel Markov Chain Monte Carlo method we call “Contour Monte Carlo”, which can be used to invert systems with a large number of parameters. We demonstrate the utility of this method by inverting a range of frequently-used deterministic models of biological systems, including the logistic growth equation, the Michaelis-Menten equation, and an SIR model of disease transmission with nine input parameters. We argue that the simplicity of our approach means it is amenable to a large class of problems of practical significance and, more generally, provides a probabilistic framework for understanding the inversion of deterministic models.

**Author summary:** Mathematical models of complex systems are constructed to provide insight into their underlying functioning. Statistical inversion can probe the often unobserved processes underlying biological systems, by proceeding from a given distribution of a model’s outputs (the aggregate “effects”) to a distribution over input parameters (the constituent “causes”). The process of inversion is well-defined for systems involving randomness and can be described by Bayesian inference. The inversion of a deterministic system, however, cannot be performed by the standard Bayesian approach. We develop a conceptual framework that describes the inversion of deterministic systems with fewer outputs than input parameters. Like Bayesian inference, our approach uses probability distributions to describe the uncertainty over inputs and outputs, and requires a prior input distribution to ensure a unique “posterior” probability distribution over inputs. We describe a computational Monte Carlo method that allows efficient sampling from the posterior distribution even as the dimension of the input parameter space grows. This is a two-step process where we first estimate a “contour volume density” associated with each output value which is then used to define a sampling algorithm that yields the requisite input distribution asymptotically. Our approach is simple, broadly applicable and could be widely adopted.

## 3 Introduction

Mathematical models have played an important role in understanding a wide range of biological phenomena, including the structure and function of the heart [1, 2], the growth of tumours [3,4], epidemics of infectious diseases [5], and population dynamics in conservation areas [6,7]. In all of these models, the variables of interest (the “outputs”) are mediated by input parameters, which, when varied, can produce a multitude of system behaviours. Whilst these parameters take algebraic form, they should (ideally) be relatable to biological processes. Determining the sets of inputs which maintain normal functioning or, alternatively, cause the system to produce unfavourable outcomes are crucial for a range of applications. Since biological systems are typically highly nonlinear [8], the sensitivity of outputs to changes in the input processes can be difficult to determine without recourse to model analysis or simulation. A forward sensitivity analysis can determine the probability distribution of the outputs of a model based on the probability distribution of model inputs. A so-called inverse sensitivity analysis determines the probability distribution of model inputs which produces a known distribution of outputs.

While forward and inverse sensitivity analyses are important concepts in all areas of science and technology, these issues are particularly acute in mathematical and computational biology. Consider first a simple physical system, the laminar flow of a Newtonian fluid which is governed by the Navier-Stokes equations, a classical and universally accepted system of partial differential equations based on fundamental physical principles and simplifying assumptions. The Navier-Stokes equations include a small number of parameters (e.g., viscosity, surface tension) that can be independently measured with high precision and laboratory-scale experiments can be conducted in well-controlled environments with a high degree of reproducibility. Modelling in the biological sciences by contrast, is currently in a rapid state of development, many models are highly parametrized, and their parameters cannot be independently measured but must be inferred from the output of the model. Even in situations when the mathematical model is well established, such as the Hodgkin-Huxley equations, some parameters may be well characterized from experimental observations, while others are not. A recent study of the Hodgkin-Huxley equations [9] indicates that sufficient membrane potential data can provide a reasonable fit to the three major conductance parameters. However, no matter how complete and accurate the membrane potential data, the time constants associated with the rate parameters in the model are practically unidentifiable. Whereas parameter estimation seeks a single value for model parameters that could produce a set of output values, the corresponding inverse sensitivity problem seeks the *distribution* of model parameters that is compatible with an observed output distribution. For example, in the Hodgkin-Huxley system, an inverse sensitivity analysis can determine how variable the unobserved rate parameters can be whilst having no real effect on the peak-to-trough voltage.

For a large class of models of practical interest in biology, including ordinary differential equations (ODEs) and partial differential equations (PDEs), analytic methods for inverse sensitivity analysis are unavailable. Further it is common for the dimension of the inputs to exceed that of the outputs. For deterministic systems, this means that there exist non-singular sets of inputs that can yield a given output and that any feasible output distribution may be caused by any member of a collection of possible input distributions. In Bayesian inference of stochastic systems, prior distributions are specified for parameters of the likelihood ensuring a unique input distribution for a given set of output data. For deterministic models, however, there is no uncertainty in the input-to-output map, meaning standard Bayesian techniques cannot be used without introducing an artificial error distribution. Here we describe a conceptual framework to understand the inversion of deterministic systems. We then describe a novel Markov Chain Monte Carlo method we call “Contour Monte Carlo” and use this approach to invert a number of systems, including the logistic growth equation, the Michaelis-Menten equation, and an SIR model of disease transmission. In contrast to methods that use a grid to discretize the parameter domain to determine a numerical output-to-input map [10], our method uses stochastic sampling to instead approximate the multiplicity of a given output value; that is, the number of inputs that map to that value. By using random sampling, the computational complexity of our approach does not grow with parameter dimensions, meaning that it can be applied to large model systems. We also argue that the simplicity of our approach means it is amenable to a large class of problems of practical significance.

<u>Outline of the paper</u>: In §4.1, we present the basic CMC algorithm assuming uniform priors in §4.1.1, and extend this approach to general prior distributions in §4.2. In §4.3, we examine several algebraic examples to illustrate how the CMC algorithm works in detail. CMC is then used to invert a range of examples drawn from mathematical biology in §5.1 before a discussion in §6.

## 4 Methods

### 4.1 Estimating the posterior distribution

#### 4.1.1 Uniform prior distributions

Consider a deterministic function from a vector of inputs 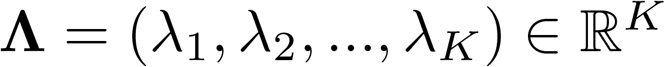 to output quantities 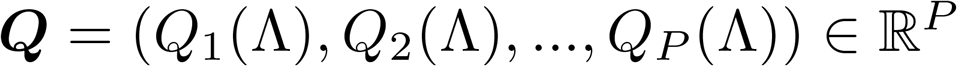, where *P* < *K*. Since the dimensionality of the input parameters exceeds that of the output quantities, each feasible output value permits a set of possible input causes. This means that without further information it is not possible to determine a single cause for an output value, unless that output value is “singular”. That which is true for individual output values is also true for output distributions. For any given output distribution, there are a collection of possible input distributions that could yield it.

The distribution that we want to estimate is a conditional density of the form *p*(**Λ**|***Q***(.), *data*), where ***Q***(.) denotes the particular form of input-to-output function used, and *data* denotes a collection of output data points. In analogy to Bayesian inference, we term this conditional density over inputs the “posterior distribution”. In what follows we implicitly assume a dependence on ***Q***(.), and omit it for brevity. To sample from this distribution efficiently, we first consider the joint distribution,

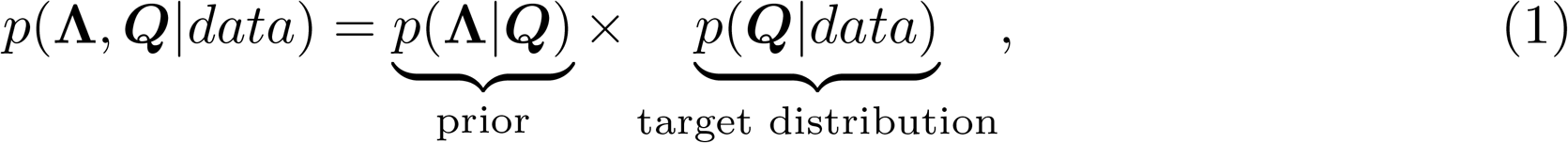

where the *prior* term does not contain any dependence on the data since it solely depends on the geometry of the input-output map, which is determined by ***Q***(.). We assume that the output data distribution has been fitted with an appropriate distribution, to yield a density value for any value of output, *p*(***Q***|*data*), which we term the “target distribution”. An alternative way to decompose the joint distribution is,

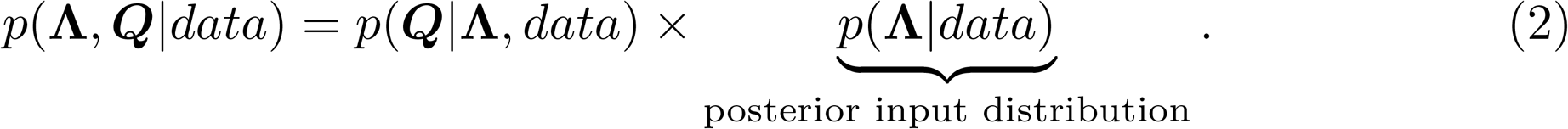

For a deterministic model *p*(***Q***|**Λ**, *data*) = *p*(***Q***|**Λ**), since ***Q*** is fully determined by the inputs **Λ**. Because of the determinacy this term becomes,

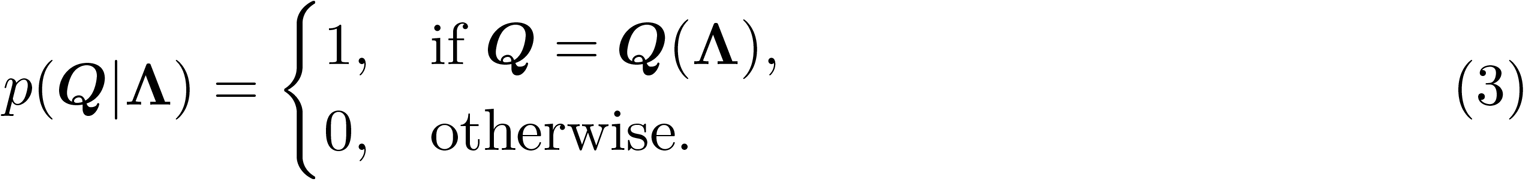

Equating the right hand side of equations (1) and (2), we obtain an expression for the posterior input distribution,

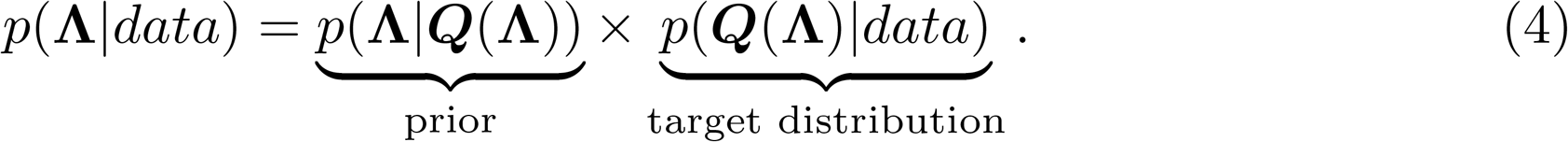

To sample from the posterior distribution specified by equation (4), we use our estimates of the prior that are obtained by independent sampling of the inputs uniformly within their bounds (we consider unbounded non-uniform priors in §4.2) to produce an estimator of the distribution,

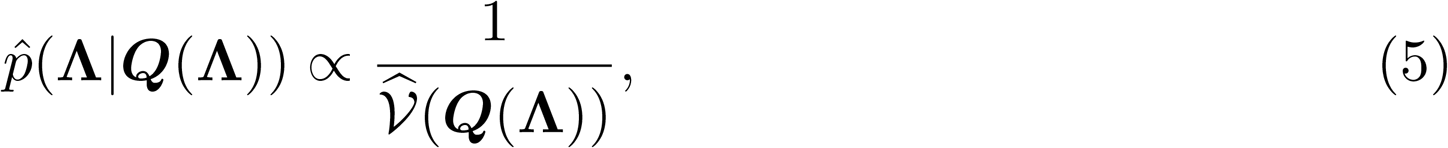

where 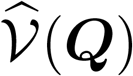 is the (potentially un-normalised) contour volume at an output value of ***Q*** - a measure of the size of the input set which maps to ***Q***. Due to the continuous nature of the output, the distribution 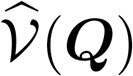 actually defines a contour volume density, which can be used to estimate the posterior input distribution,

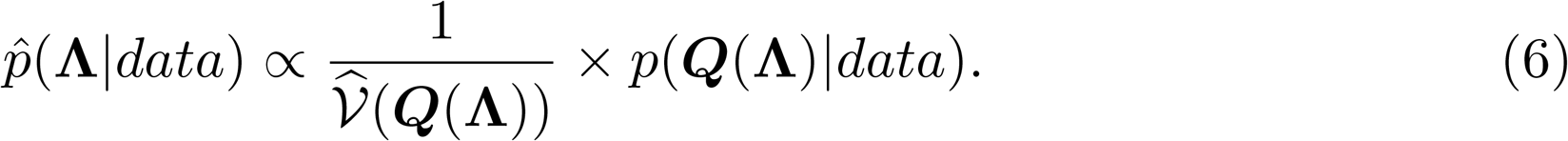

Since the above probability distribution is non-standard and potentially un-normalised, we use MCMC to sample from it, and use Random Walk Metropolis sampling [11] for this process with an acceptance probability of,

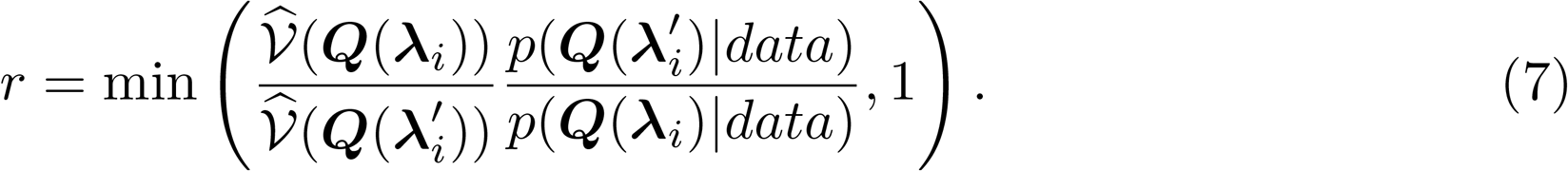

We recognise that since an approximate form of the un-normalised distribution, as well as its gradients, are known, more efficient forms of MCMC algorithms such as adaptive Metropolis [12], or Hamiltonian Monte Carlo (HMC) [13] could also be used.

To summarise, our “Contour Monte Carlo” (CMC) algorithm is composed of two distinct phases (see Algorithm 1 for more detail): first, sample the input parameters to yield estimates of the contour volume density, 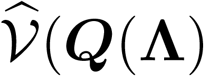; second, use 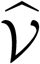 to specify a prior that distributes probability mass uniformly within a contour volume which, along with a target output distribution, determines a posterior input distribution. This distribution is then sampled from using an MCMC method.

##### Algorithm 1

Pseudocode for the Contour Monte Carlo algorithm for parameters with uniform probability distributions on bounded domains (Λ). (Parameters that may be vectors are shown in bold.)

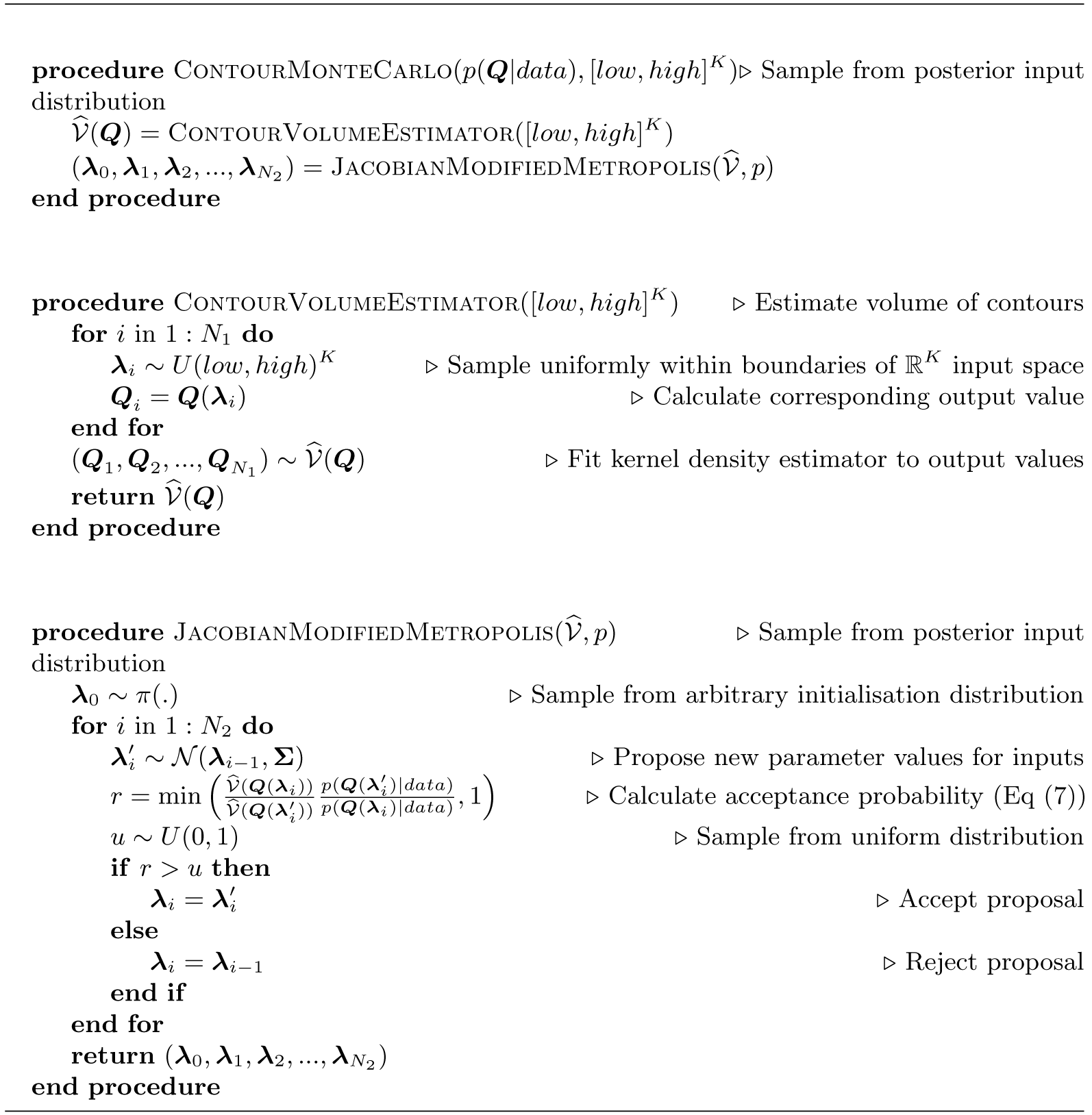

Since we are reconstructing the posterior input distribution by MCMC sampling, it is crucial to determine when our sampling distribution has converged. To do so, we use the same convergence diagnostic criteria that are used in applied Bayesian inference. We run a number of Markov Chains in parallel and compare their within-chain variance to that between the chains by calculating 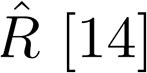 [14], and use a convergence threshold of 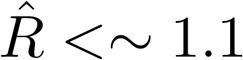 on all input parameters. Once convergence is achieved, we discard an initial portion of the chains as “warm-up” [11, 15] and use the rest to form our posterior distribution.

### 4.2 General prior distributions

The type of uniform prior we have described thus far appears sensible since it allocates probability mass in exact accordance with the geometry of the input-output map. In the absence of further information it seems reasonable to suppose that all inputs which produce a given output value are equally likely causes of it, and Algorithm 1 describes how to sample from the resultant posterior input distribution. It is unclear, however, how to use our existing CMC algorithm to handle unbounded prior spaces, where uniform prior distributions cannot be specified. We now extend the framework described in §4.1.1 to handle arbitrary prior distributions. We start with the expression for the posterior distribution,

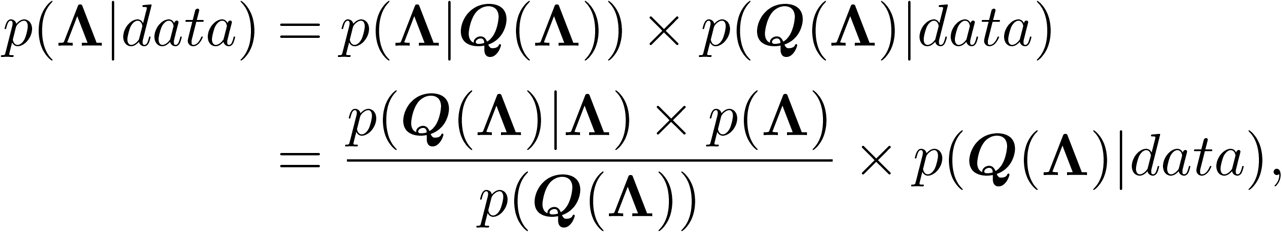

where we have used Bayes’ rule to rewrite the conditional prior term. We then recognise that for a deterministic model *p*(***Q***(**Λ**)|**Λ**) = 1, resulting in the following general expression for the posterior distribution,

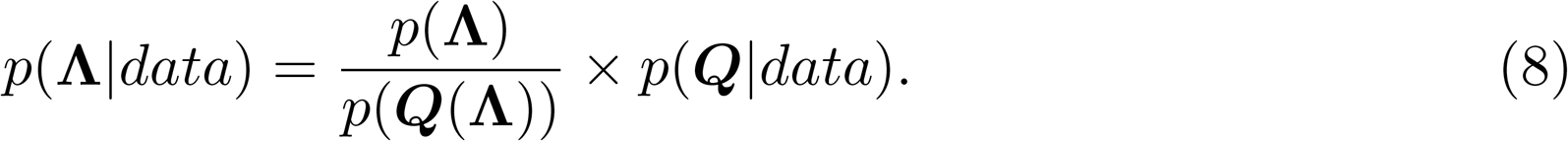

If the parameter bounds are non-infinite and consist of a total volume *V* in input space, and we suppose that all values of **Λ** are equally likely *a priori*, then the above becomes,

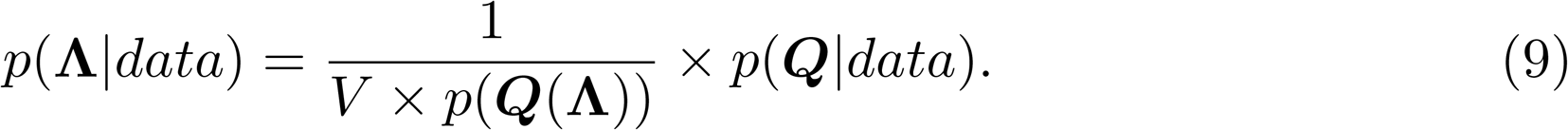

However if all parameter values are equally probable then *p*(***Q***(**Λ**)) is simply the proportion of input space defined by the set {Λ′: *Q*(Λ′) = *Q*(Λ)}. Multiplying this by the overall volume of input space yields the volume of input space occupied by this set, 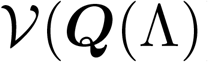, and we recapitulate the expression for the posterior that we approximated in equation (6). We thus see that the assumption of a uniform prior within a contour volume is equivalent to the assumption of an uniform prior throughout the input domain.

More generally, we can imagine setting informative priors on parameters that may or may not be finitely bounded. For parameters with infinite bounds a uniform prior is improper, and we therefore resort to using valid probability distributions that have support over an infinity of values. In general, the priors that we set will not be aligned with the shape of input contours, meaning that there can be a variation in the probability density within a given contour volume. How should we handle parameter inference in this new framework? Equation (8) is the expression for the posterior in this, more general, setting. Algorithm 2 describes how to account for general prior distributions in CMC, which is then used in illustrative example 3(b) to estimate the posterior input distribution.

#### Algorithm 2

Pseudocode for the Contour Monte Carlo algorithm for parameters with general probability distributions on potentially unbounded domains (Λ). (Parameters that may be vectors are shown in bold.)

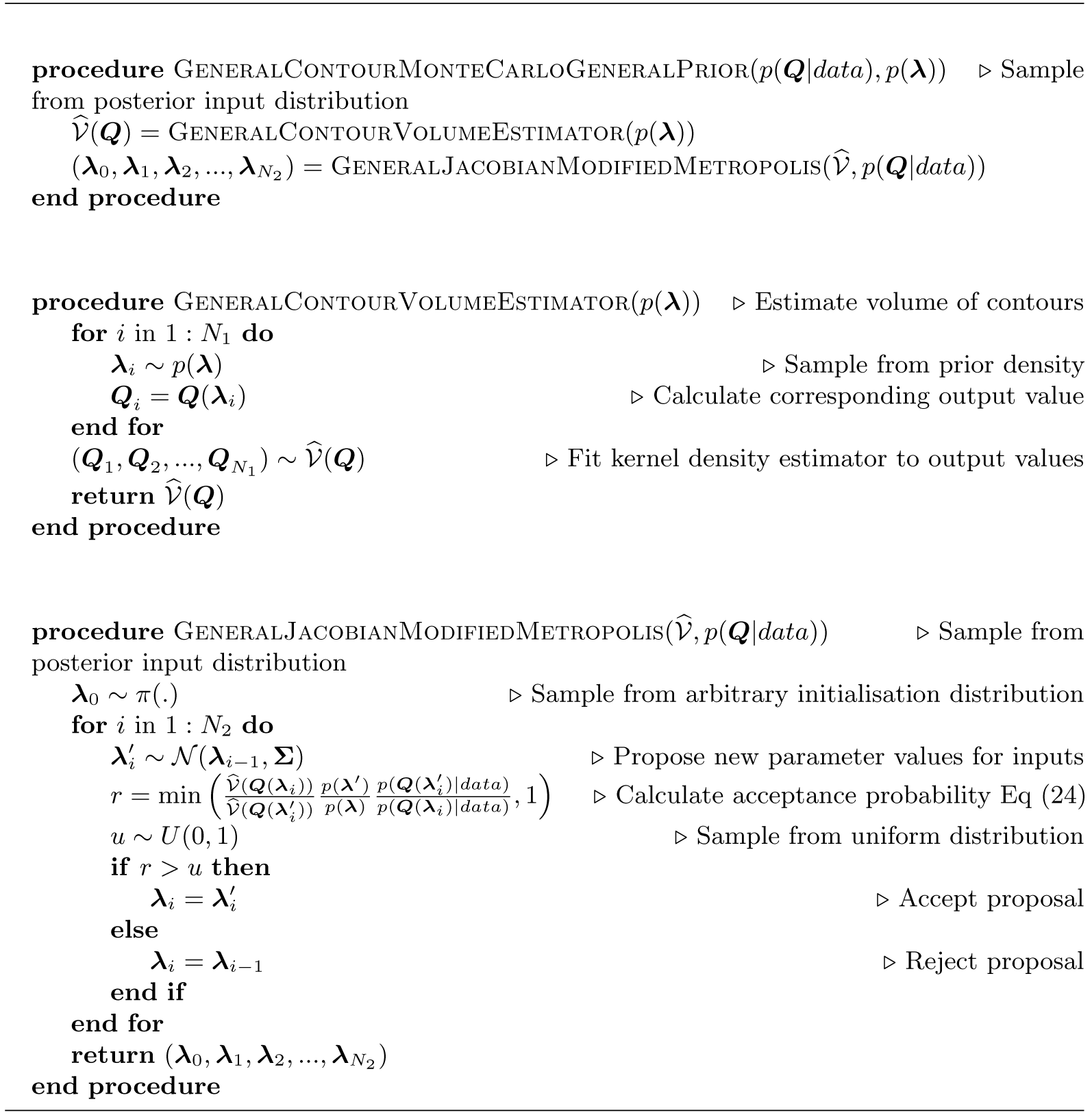

### 4.3 Illustrative examples

#### 4.3.1 One-to-one input-to-output maps

Whilst our approach is intended for systems where the number of outcomes exceeds the inputs, simpler systems with one input and one output (one-to-one) can be used to build an intuitive understanding of the method. Here the usual Jacobian of a coordinate transformation plays the role of the prior term, *p*(**Λ**|***Q***(**Λ**)).

**Illustrative example #1**

Consider an input (*λ*) to output (*Q*) map of the form,

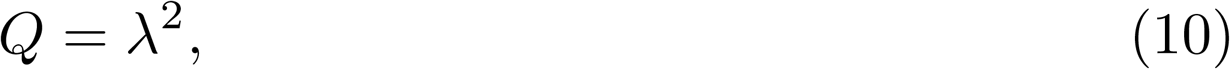

where *λ* ∈ [0, *1*]. Suppose that we seek a distribution over inputs *p*(*λ*) such that the corresponding distribution over outputs follows *Q*(*λ*) ~ *beta*(2, 2) (see the black lines in the right hand column of Figure 1). A straightforward way to generate samples from *p*(*λ*) is to first sample *Q_i_* ~ *beta*(2, 2) then use the inverse map to yield samples of 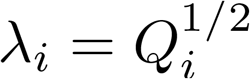. For most interesting models in mathematical biology however, including ODEs and PDEs, an inverse map cannot be calculated or does not exist. As such, we aim to generate samples from *p*(*λ*) using only the forward map.

**Figure 1.**
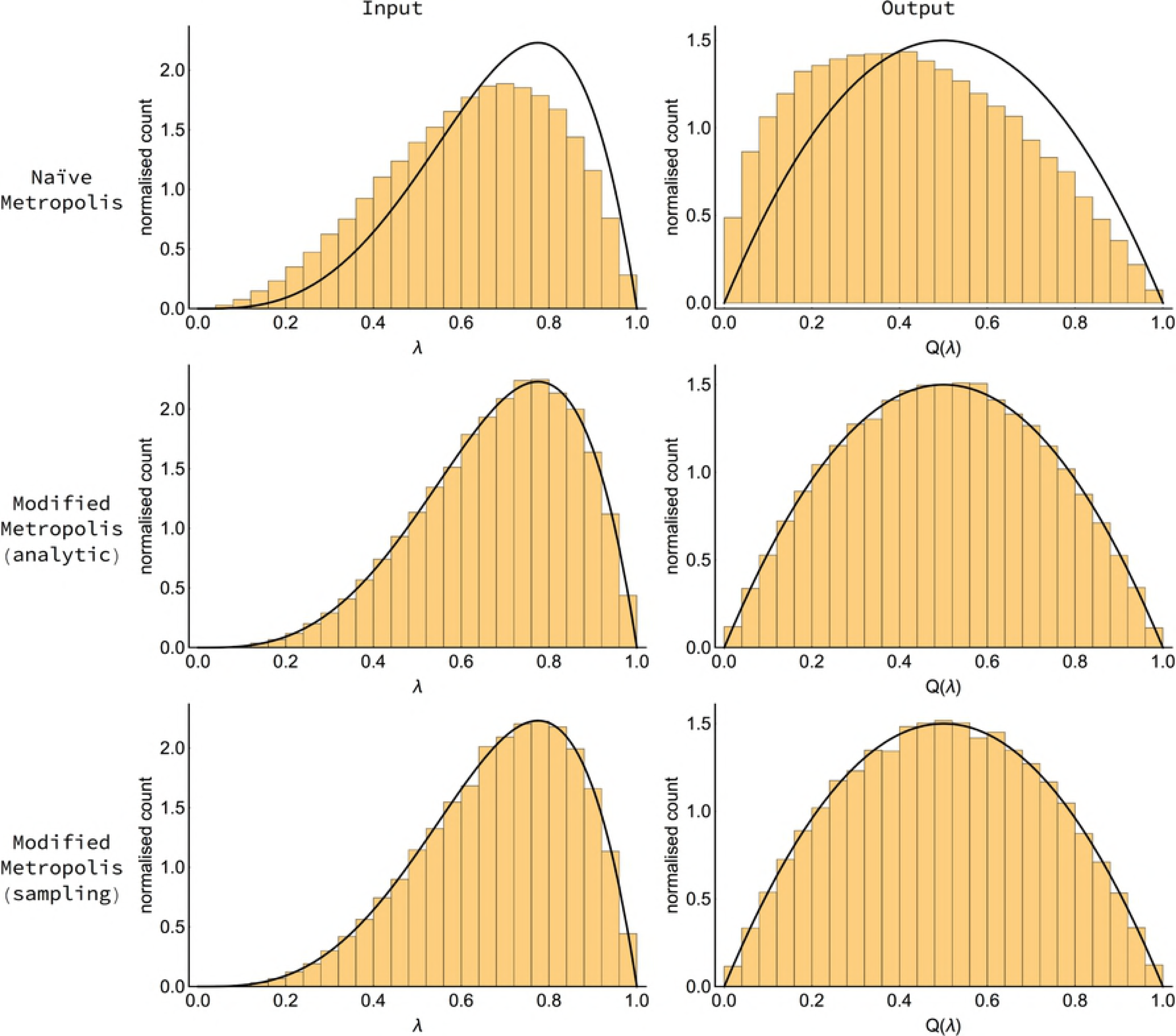
Illustrative example #1: Estimated posterior input distributions (left column) and corresponding output distributions (right column) for naive Metropolis (top row), modified Metropolis using the analytic Jacobian (middle row) and sampling-based Modified Metropolis (bottom row). Target densities are indicated by black lines. For the analytic case, the Jacobian used corresponds to equation (12). In each case, 316,000 Metropolis samples across 8 Markov chains (40,000 iterations on each chain, with the first 500 samples discarded as warm-up) were used to produce the histograms. For the sampling-based Modified Metropolis, the Jacobian transformations were estimated as indicated in the caption to Figure 2, and are shown as the blue-dashed line in Figure 2B.

It may appear that a valid approach to generate input samples from *p*(*λ*) is to use a Markov chain Monte Carlo (MCMC) algorithm like random walk Metropolis [16], where proposed *λ*′ are accepted as samples if,

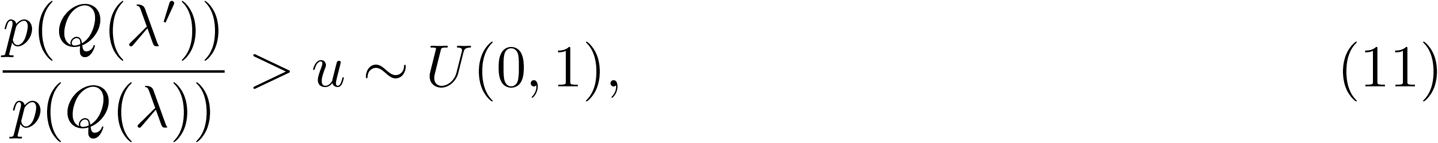

where *p*(*Q*(*λ*)) is the target density (here a *beta*(2, 2) distribution) evaluated at *Q* = *λ*^2^. This approach, however, results in an input distribution (top-left panel in Figure 1) which, when transformed, does not recapture the target density (top-right panel in Figure 1). The reason for this bias is that it neglects to include a Jacobian term in the density which accounts for the nonlinear change of measure in going from input to output space. If this Jacobian term is included in the Metropolis accept-reject rule,

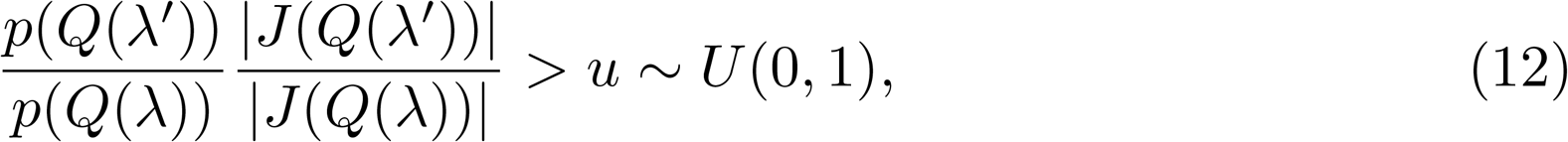

where, in this case, 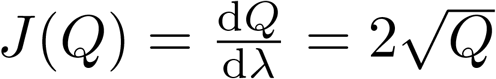, then this results in an input distribution (middle-left panel in Figure 1) which, when transformed, results in an output distribution that corresponds to the target (middle-right panel in Figure 1).

Indeed, in this simple example, it is possible to exactly determine the input distribution that results in a *beta*(2, 2) target. This is given by a Jacobian transformation multiplied by the target density,

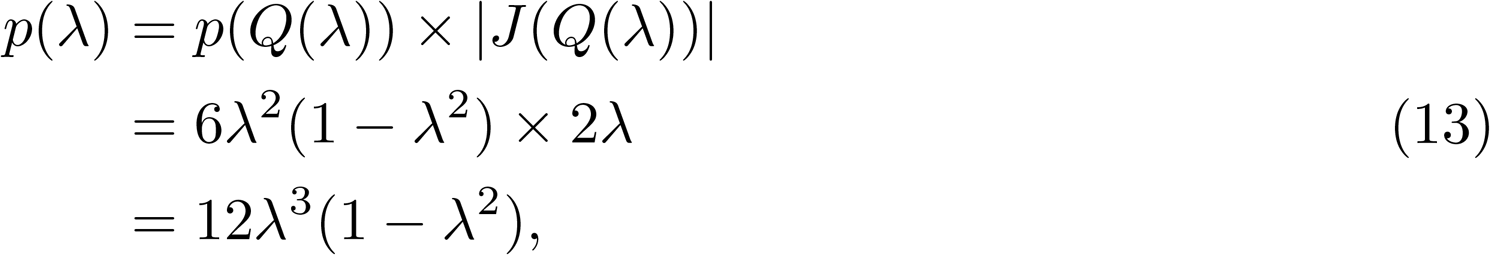

which is shown by the black lines in the left hand column of Figure 1.

##### Estimating the Jacobian transformations by sampling

Whilst the Jacobian may be calculated analytically and the marginalization performed for simple input to output maps, for maps of modest difficulty, this term cannot typically be analytically derived. We now illustrate how approximate Jacobian transformations can be obtained by sampling. To do so, we first sample uniformly from the bounds of *λ* (Figure 2A) which, when transformed by equation (10), yields a sampling distribution for *Q* (Figure 2B). By fitting a Gaussian kernel density estimator to this sampling distribution (dashed blue lines in Figure 2B), we can well approximate the analytic inverse Jacobian density (grey line) with a modest number of samples. This, hence, allows us to use the Jacobian-corrected Metropolis sampler defined by the acceptance ratio of equation (12) to generate samples from the input distribution *p*(*λ*).

**Figure 2.**
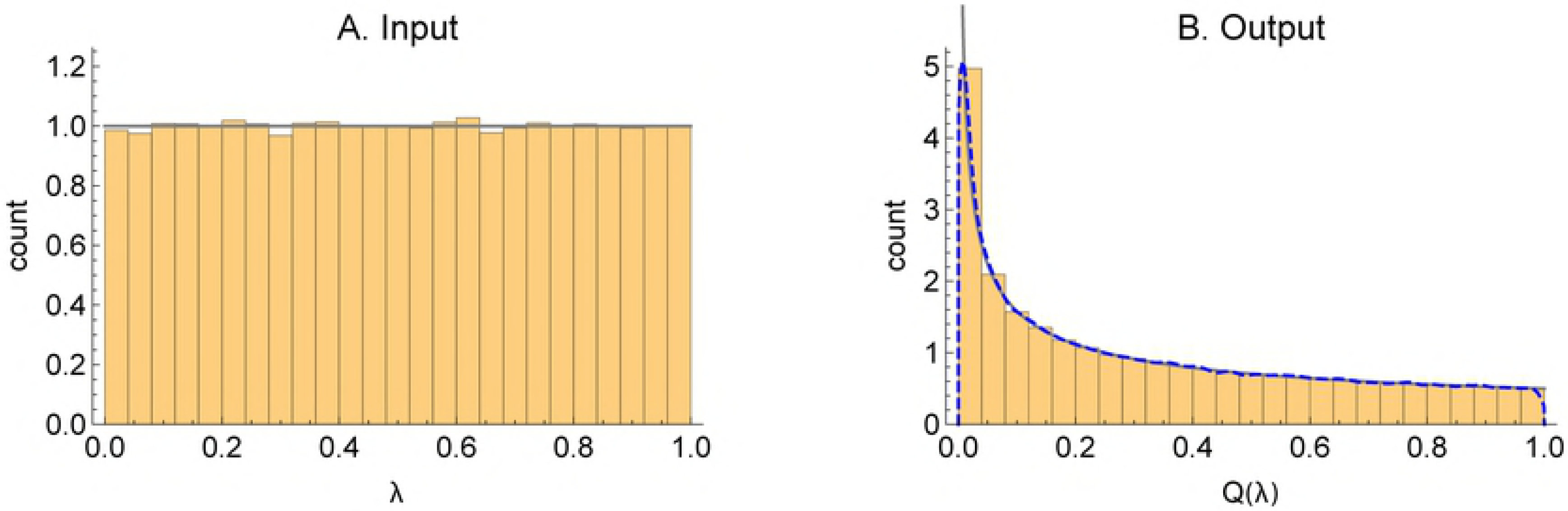
Illustrative example #1: (A) Uniform sampling of input over [0, 1]. (B) Corresponding output under map defined by equation (10). Histograms were constructed using 100,000 random samples. The grey curves indicate the exact sampling distributions. The blue-dotted line in panel B shows the Gaussian kernel density estimate of the sampling distribution using a bandwidth of 0.01.

Once the inverse Jacobian transformation has been estimated, we simply replace the actual Jacobian in our Metropolis-based rule given by equation (12), with the sample-based estimates 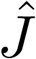,

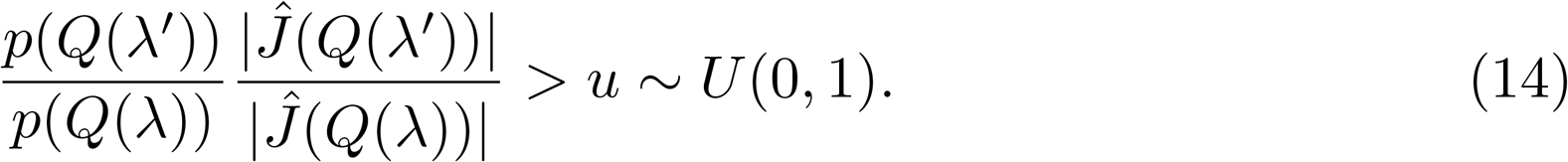

Our approach, therefore, consists of two distinct steps: first, independent sampling from the prior space (which is currently assumed to be bounded but, in §4.2, this is relaxed) and transformation of these inputs to form a sample of outputs, followed by the fitting of an empirical density estimator to the output samples; second, Jacobian-modified Metropolis sampling using the Jacobian transformations estimated from the first step. When the sample size of the first step approaches infinity, the Markov chain in the second step converges asymptotically to the target density. For a finite sample size for the first step, the sampling distribution of the Markov chain in the second step does not exactly converge to the target distribution due to Jensen’s inequality. For even modest sample sizes, however, we have found this bias negligible compared to other sources of uncertainty.

In the bottom row of Figure 1, we illustrate how using our two step method allows us to generate an input distribution which maps to the target.

#### 4.3.2 Many to one input-to-output maps

For many-to-one maps, no inverse map exists, meaning that a given output value (unless a singular point) corresponds to a number of input vectors. This means that a given output distribution can be produced from a variety of input distributions. Like in Bayesian inference, to reduce the set of allowable input distributions to one, we specify a prior distribution over input parameters. In contrast to Bayesian inference, this prior is specified as a conditional probability distribution, where we condition on particular output values.

A natural way to specify such a prior is to assume that the probability distribution is uniform within the set of possible inputs that generate a given output value. The prior distribution is then given by the probability density,

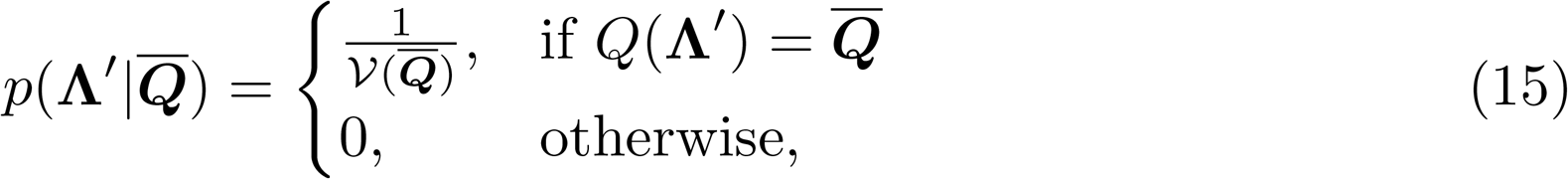

where 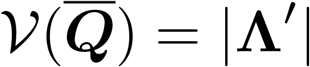 is the volume of input space where 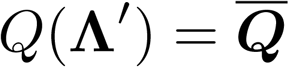. We term the size of the region of input space that produces a particular output value, its “contour volume”. Note that we use the term “volume” liberally here. For a two-dimension input space and one dimensional output space as shown in Figure 3 this corresponds to a length, although for higher dimensional examples the set can be an area, a volume or a hypervolume.

**Figure 3.**
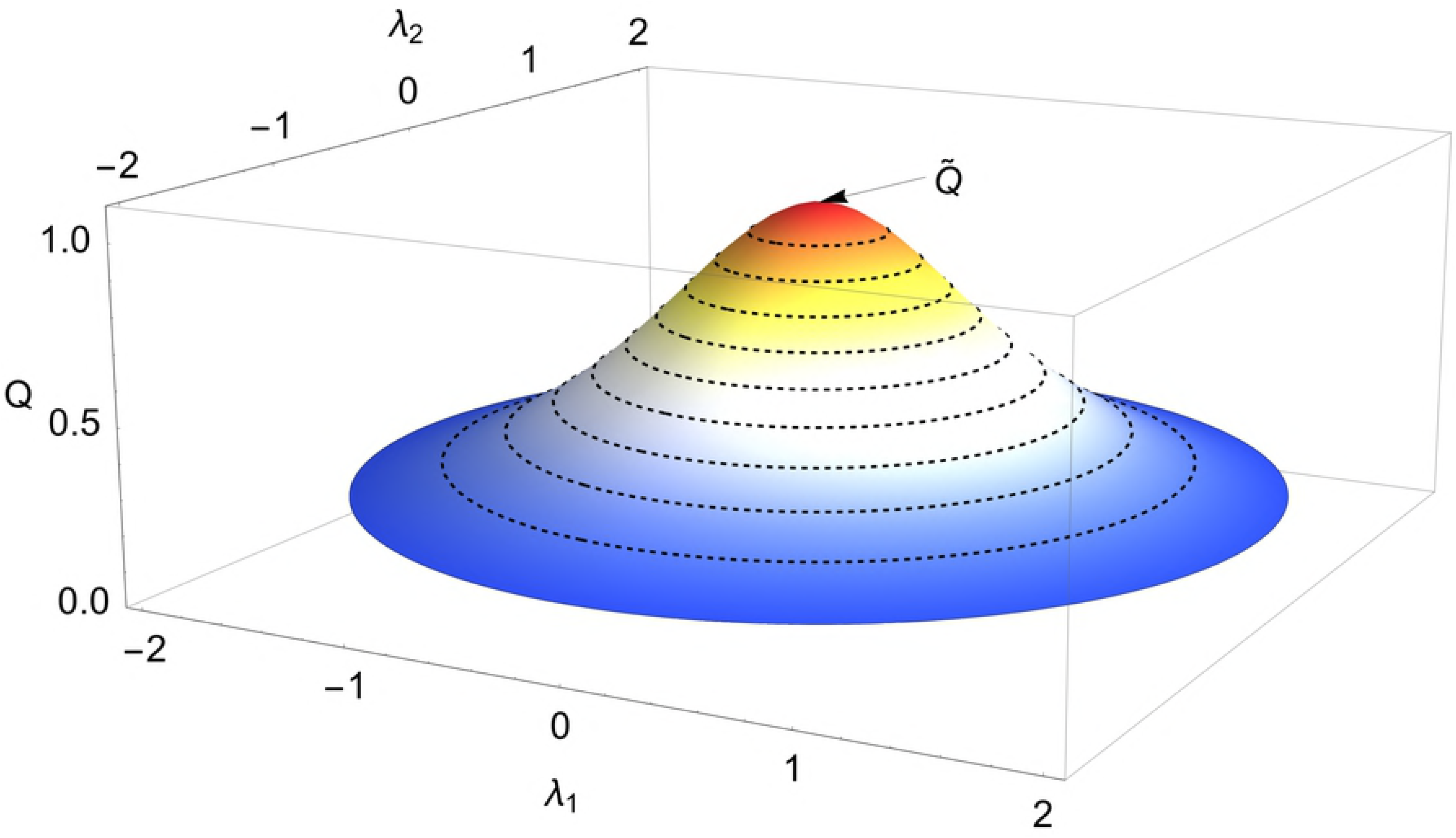
Illustrative example #2: Plot of function given by equation (16) showing contours of constant *Q* values. Here 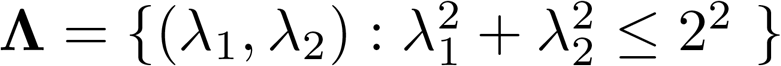.

**Illustrative example #2**

To explain how this process works we consider the case of two dimensional inputs (*λ*_1_*,λ*_2_) bounded by the region 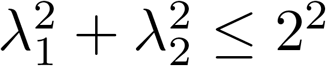, and a uni-dimensional output,

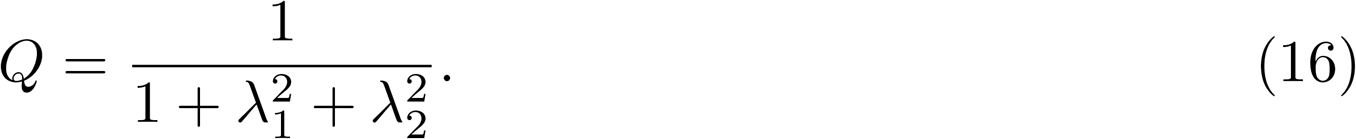

A plot of this function is shown in Figure 3, where it is observed that each value of the function 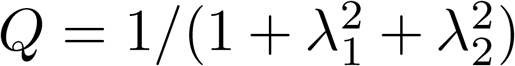 maps to any point along circular “contours” (dashed lines) in input space (*λ*_1_*,λ*_2_), with the exception of singular point 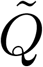.

We suppose that we want to produce an input distribution which results in a target output distribution which is uniform across allowed function values *Q* ∈ [0.2, 1], since the function is bounded by its extreme values at 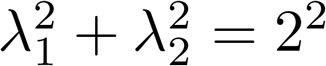 (where *Q* = 0.2), and (*λ*_1_*,λ*_2_) = (0, 0) (where *Q* = 1).

We seek an analytic expression for the input distribution which is consistent with this target distribution given a uniform prior distribution over inputs. The prior distribution which is uniform over inputs can be transformed from Cartesian 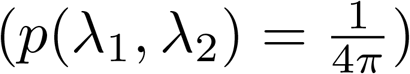 to polar coordinate form,

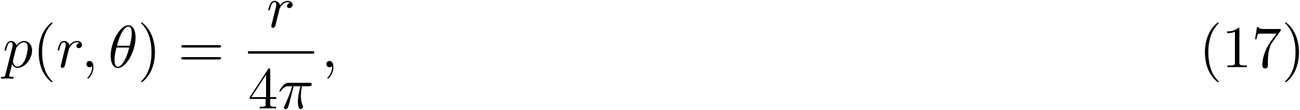

where 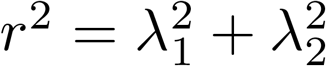 and *θ* = arctan(*λ*_2_/*λ*_1_). When marginalising over *θ* the above expression yields *p*(*r*) = *r*/2. Since we can rewrite *Q* = 1/(1 + *r*^2^), a one-dimensional Jacobian transformation yields the contour volume distribution pertaining to *p*(*r*),

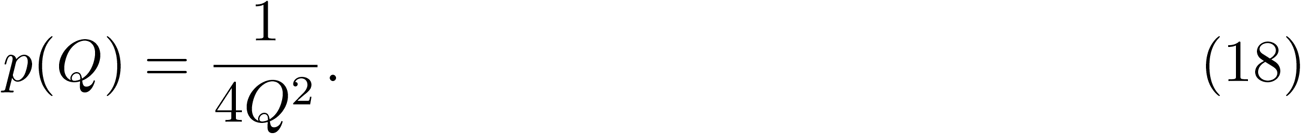

*Note:* The “Jacobian” associated a nonlinear change in measure for a mapping from 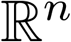 to 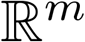 where *m* < *n*, is constructed in Theorem 2.1.1 of [17].

For this example, we can analytically calculate the prior term in equation (4) using equation (15) and the normalizing constant equal to the area of the domain Ω ( = 4*π*),

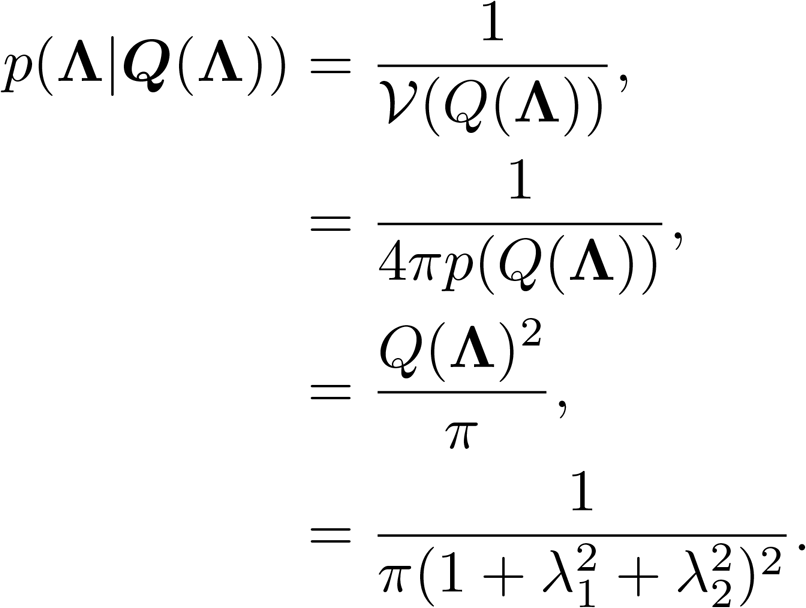

Then using Eq (4), and targeting a uniform output distribution across the range of *Q*, *p*(*Q*|*data*) = 1/(4/5) = 5/4, we calculate an analytic form of the posterior input distribution,

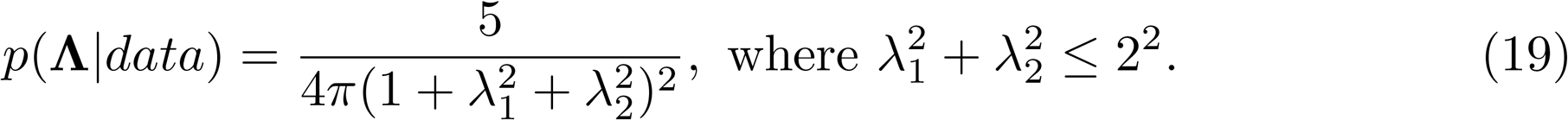

This can be alternatively expressed in polar coordinates as,

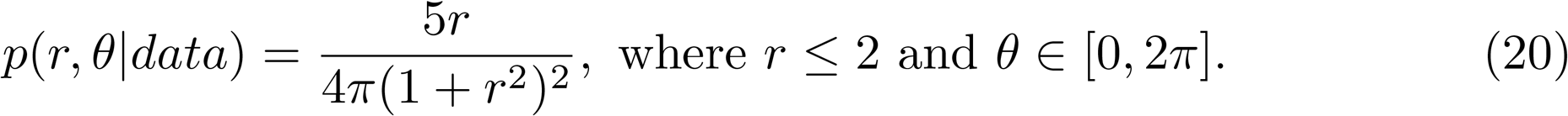

If we marginalise *θ* from equation (20), we obtain the marginal posterior distribution for *r*,

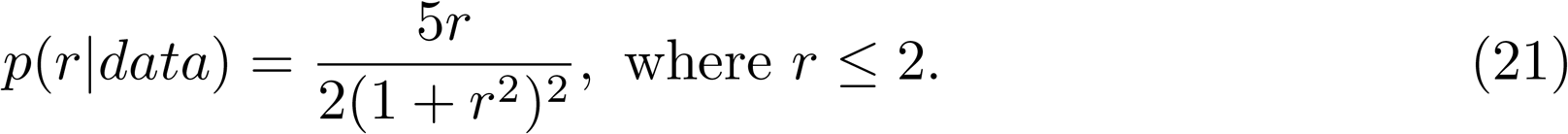

In most applied cases, analytic calculation of the contour volume distribution, as is given by equation (19) in this example, is not possible, and we must instead approximate it by sampling. We now illustrate this process for the current example. To start, we idealise the problem and assume the function value *Q* is constant within annuli of a given width and distance from the origin (Figure 4A). Since our function and input domain are well-behaved it is possible to analytically calculate the area of each annulus (Figure 4B).

**Figure 4.**
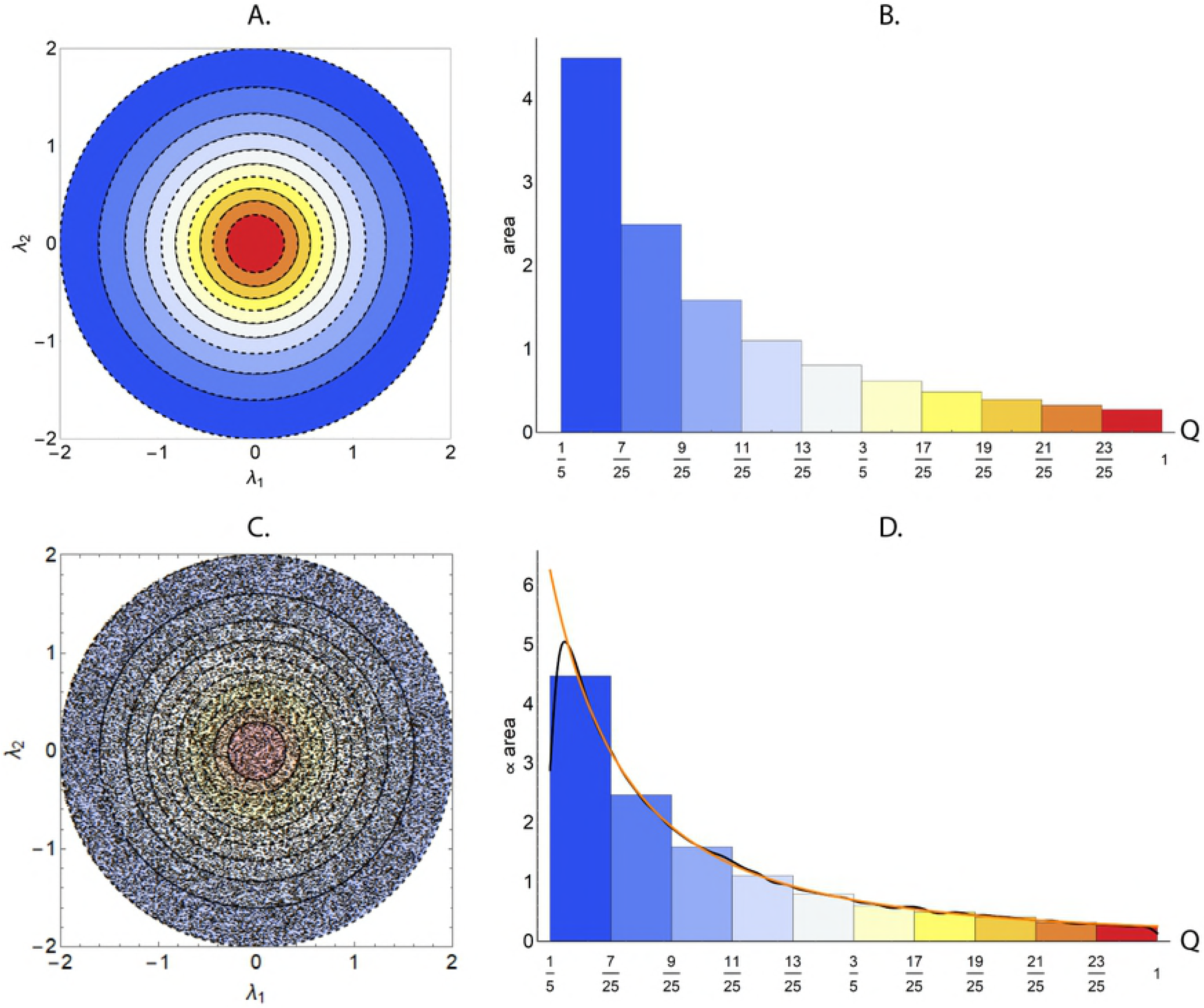
Illustrative example #2 (A) Mean values of the function given by equation (16). (B) Areas of each annulus shown in subplot A. (C) Uniform sampling of the input space. (D) Contour volumes estimated by the uniform sampling indicated in subplot C. Subplot C was produced using 50,000 independent samples of the input. The dashed lines indicate contours of the function. The black line in subplot D shows a kernel density model fit estimated using the un-binned output data with Gaussian kernel and a bandwidth of 0.01 (using Mathematica’s “SmoothKernelDistribution” function [18]). The orange line indicates the true contour volume distribution. Here 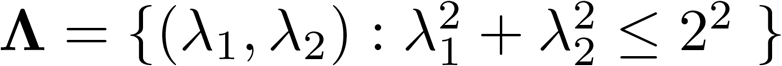.

In the above explanation, we assumed that the function value was constant within a given annulus, however we recognise that the continuous nature of the function means that the contours are actually one dimensional lines (Figure 3). Given the simple nature of this output function we can actually analytically determine the length (which we call a contour volume) of each of these lines. In our example, the output space is a foliation of these one-dimensional lines, whose non-uniform spacing determines a density of contour volumes (roughly, the area of input space that maps to an infinitesimal range of output). In Algorithm 1, we estimate the contour volume distribution by sampling uniformly from the input space (Figure 4C), and for each set of inputs 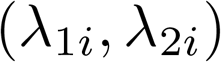, we calculate a corresponding *Q_i_*. To approximate *p*(*Q*) in equation (18) we then fit a kernel density model to (*Q*_1_*,Q*_2_, …, *Q_N_*), where *N* is the number of input samples (the black line in Figure 4D). After relatively few input samples, the estimated contour volume distribution well approximates the true distribution of equation (18) (the orange line in Figure 4D).

Note that the contour volume distribution *p*(*Q*) is equivalent to the two-to-one inverse Jacobian transformation for this problem. Specifically, Figure 4D is analogous to Figure 2B, except for a two-dimensional input example. In all cases where the input spaces are bounded and the input priors are uniform, what we call the “contour volume distribution” is equivalent to the density defined by the inverse Jacobian transformation.

We compare the estimated marginal posterior distribution for *r* that results from the CMC algorithm with the analytic result of equation (21). In the first phase of CMC, we estimate the contour volume for any output value using a kernel density estimator fit to the function evaluations from uniformly sampled inputs (Figure 4D), and use this in a second phase targeting a uniform output space (within bounds). In this case, our estimated posterior density is of the simple form,

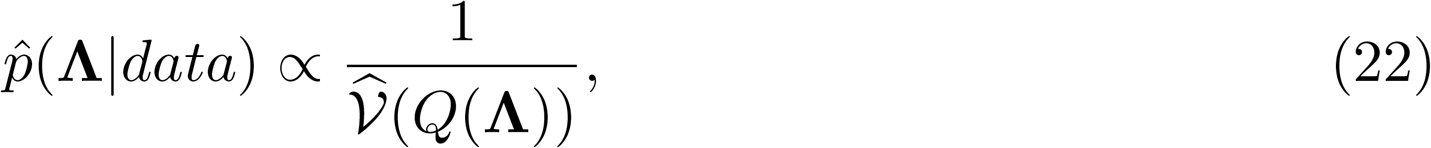

in other words we step in inverse proportion to the estimated contour volumes. This results in a marginal posterior density for inputs which is concentrated towards the centre of the input domain (Figure 5A), which results in a uni-dimensional distribution for 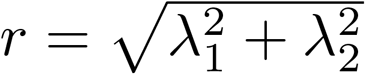 which is in agreement with the analytic result of equation (21) (Figure 5B). A benefit of our algorithm is that we can validate it by comparing the marginal output posterior with the target. Since in this case our output distribution looks approximately uniform (Figure 5C), it appears that the algorithm is working correctly.

**Figure 5.**
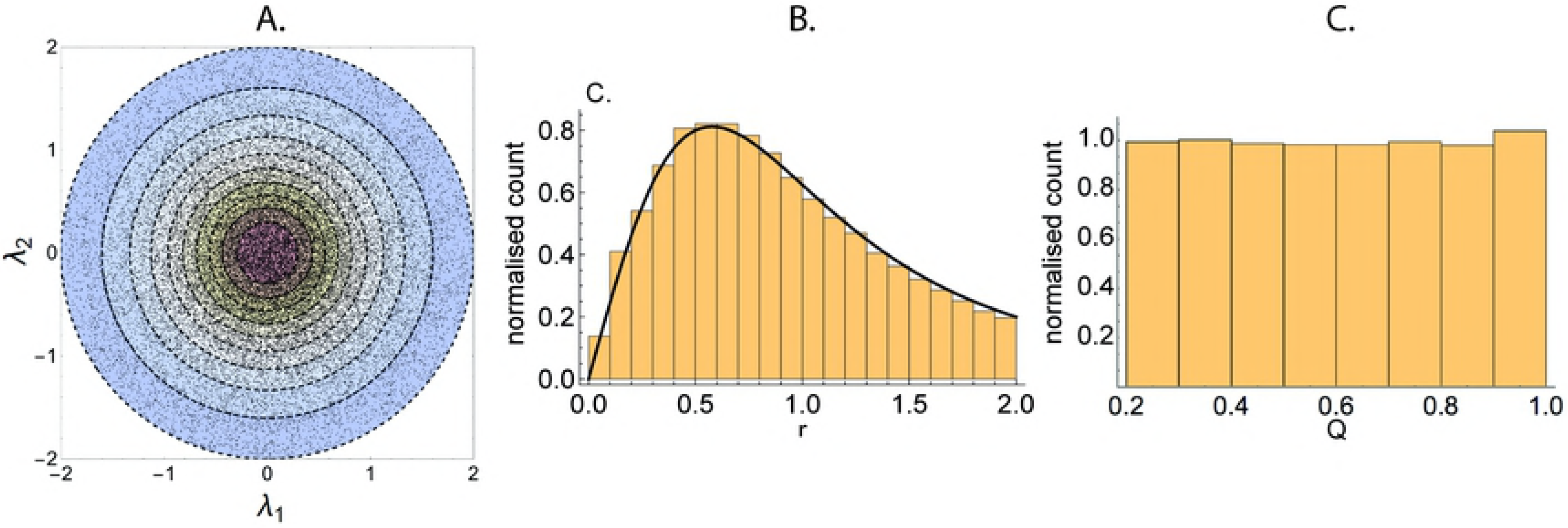
Illustrative example #2: (A) Joint posterior distributions over inputs (*λ*_1_,*λ*_2_). (B) Posterior distribution in terms of 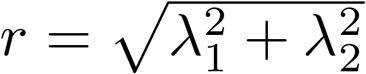 versus the analytic result (solid line). (C) Posterior distribution over outputs. The analytic result is given by equation (21). The kernel density estimator for the volume of contours is shown in Figure 4D and resulted from 50,000 independent samples from a uniform distribution over input space. The MCMC was performed using the Metropolis algorithm with toroidal boundaries and consisted of 100,000 samples on each of 12 chains. Here 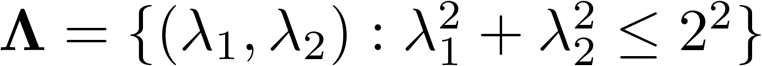.

In Illustrative example 2, a sampler using a *Q*(Λ)|*data* ~ *u*(0.2, 1) distribution over output values will accept all input samples, and hence samples uniformly in input space (Figure 4C). The result is a sampling distribution of inputs with more points towards the boundary of the input domain. Those boundary values of the domain have a lower output value and hence the output distribution is skewed towards those function values (Figure 4D). To correct for the over-representation of points with low function values, we must correct for their longer lengths, which we do by estimating 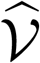 and using the inverse of this as a prior.

A more complex test problem comprised of a pair of nonlinear maps (two outputs) with three parameters that was introduced in [10] is analysed in S3–A nonlinear map.

#### 4.3.3 Informative prior distributions

We consider the output generated by a non-uniform distribution of parameters and a nonlinear map which we seek to recover assuming uniform and nonuniform prior distributions.

**Illustrative example #3**

Consider a bivariate normal as an input distribution with mean and covariance matrix (Figure 6A),

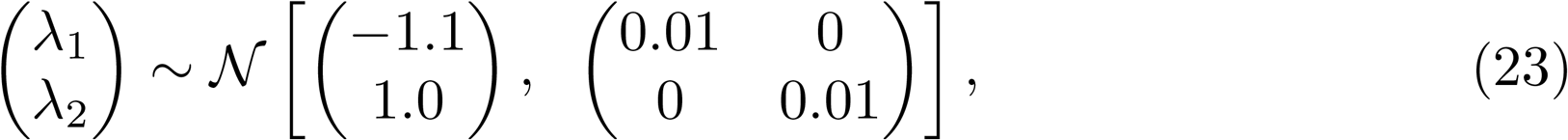

**Figure 6.**
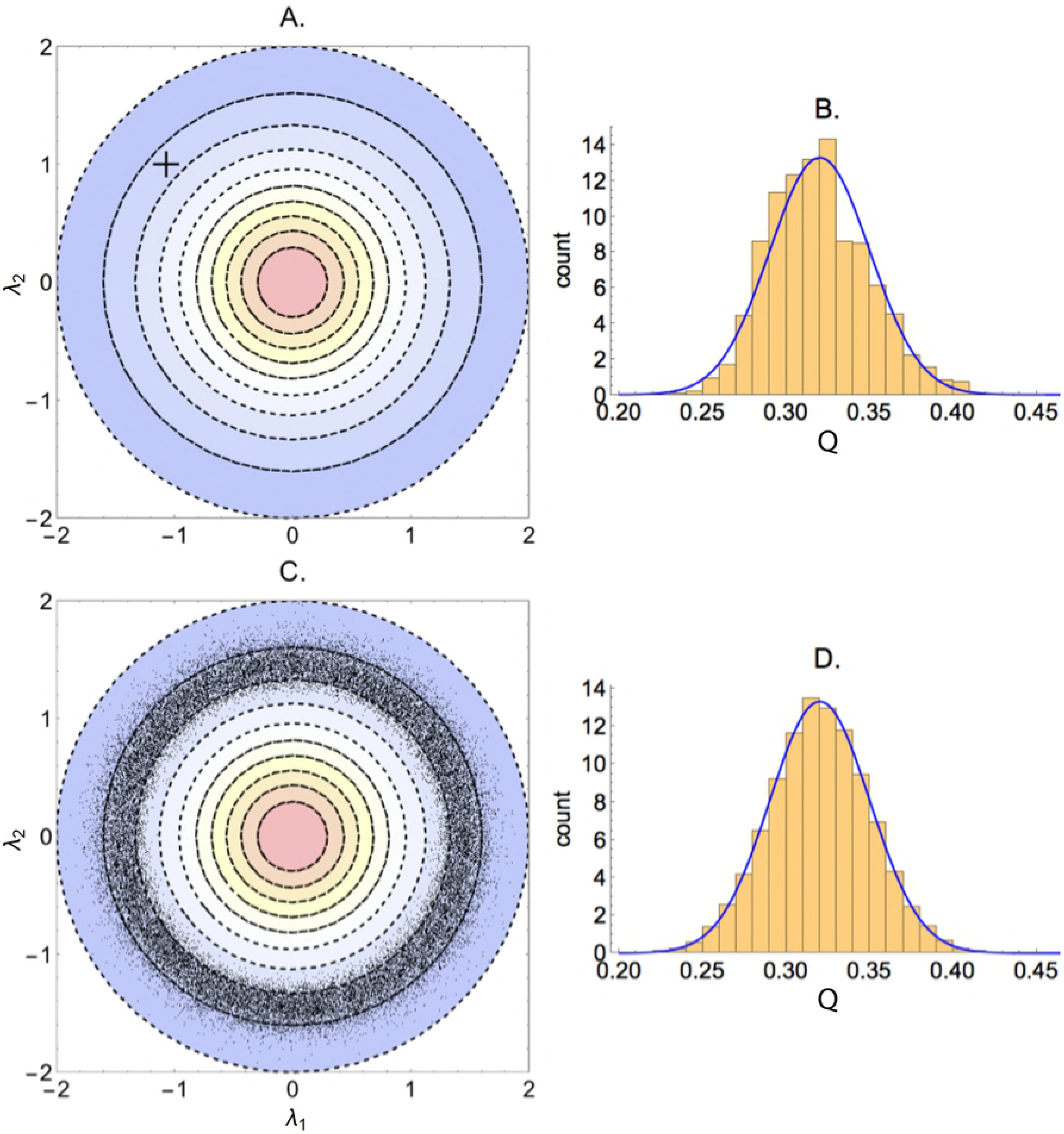
Illustrative example #3(a): (A) The cross indicates the mean and standard deviations of a bivariate normal distribution. (B) Output distribution generated by the input distribution indicated in subplot A. The blue curve indicates the fit with a normal distribution. (C) The resultant MCMC samples from the posterior. (D) The output distribution corresponding to samples in subplot C (orange histogram) and target distribution (blue curve). The kernel density estimator for the volume of contours is shown in Figure 4D and resulted from 50,000 independent samples from a uniform distribution over input space. The MCMC was performed using the Metropolis algorithm with toroidal boundaries and consisted of 10,000 samples on each of 10 chains (with the first 500 observations discarded as warm-up). Here 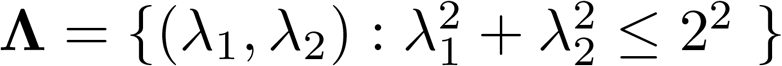.

We apply the function specified by equation (16) to a large number of independent samples from the above distribution, and we fit a normal distribution to the output distribution (Figure 6B).

##### (a) Uniform prior distribution

We then use the output distribution in Figure 6B as a target for CMC. The resultant posterior distribution is concentrated among points that lie within the same annulus in which the bulk of the probability mass for the original input distribution lies (Figure 6C). This is because our prior distribution gives equal weight to all points within a given contour. Since the function values in the annulus are similar to those obtained from using the input distribution, we obtain a posterior probability mass that is distributed uniformly around the annulus.

Note that the posterior distribution is *not* the same as the input distribution. This example demonstrates an important characteristic of deterministic models where the input dimension exceeds that of the outputs: a given input distribution induces an output distribution which, when inverted, produces an input distribution that is not necessarily the same as its cause. Information is lost when transforming from the inputs to the outputs that cannot necessarily be regained by inverting the output map.

##### (b) Normal prior distribution

We now change the priors such that they coincide with the input distribution given in equation (23). We sample from this prior distribution (Figure 7A) and use each (*λ*_1_*_i_,λ*_2_*_i_*) pair as inputs to *Q*(*λ*_1_*_i_,λ*_2_*_i_*). We then obtain an approximate output distribution by fitting a kernel density estimator to the resultant output values (Figure 7B). Note that because our prior distribution has changed, our prior output distribution 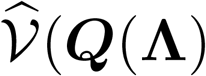 changes (compare Figure 7B with Figure 4B). We use this estimated output distribution as an input to the Metropolis stage of CMC (see Algorithm 2), where we accept an input proposal with probability,

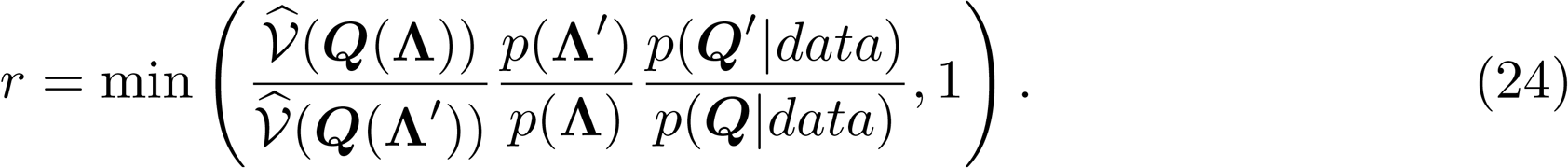

**Figure 7.**
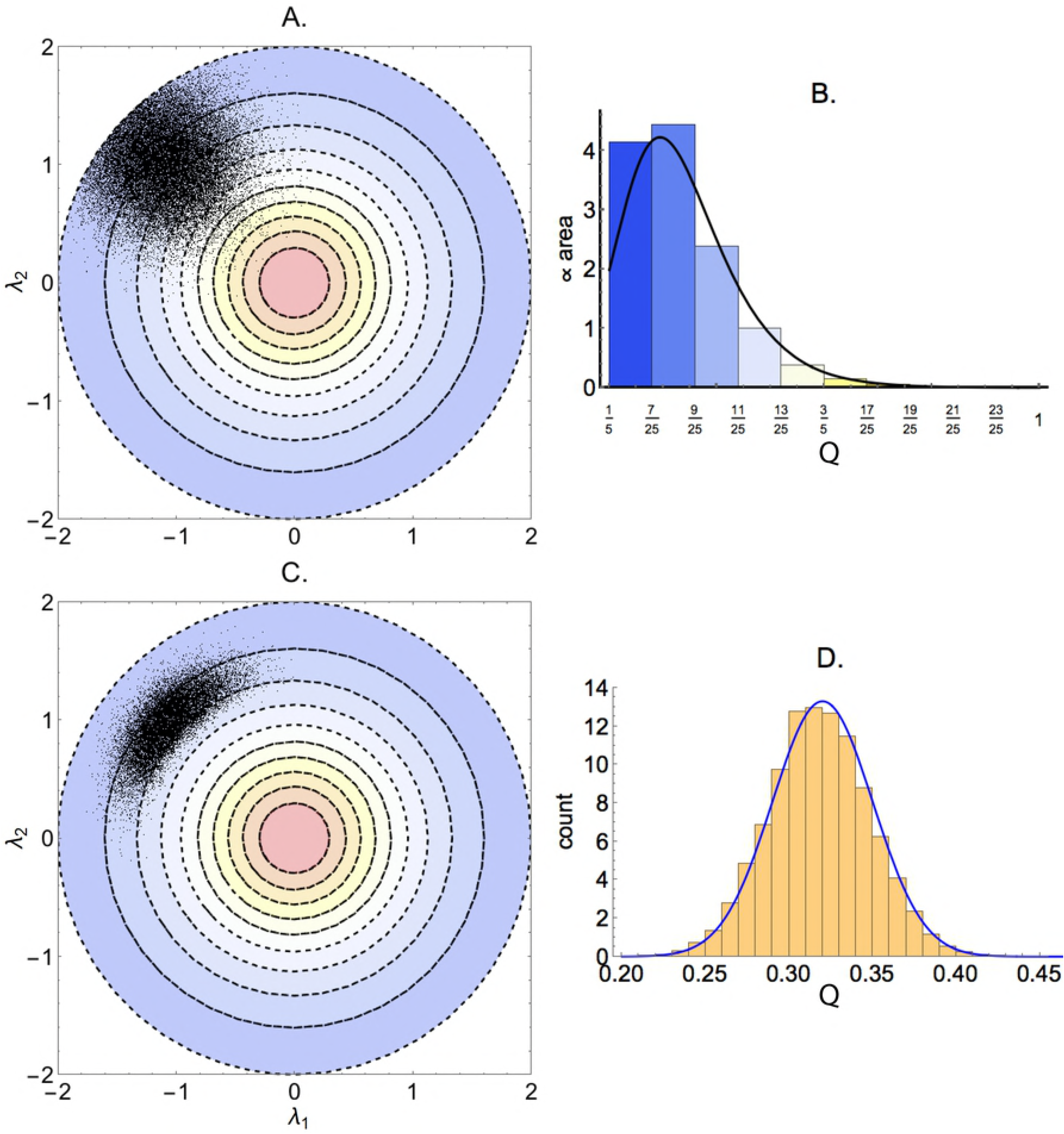
Illustrative example #3(b): (A) Samples from the prior distribution specified by equation (23). (B) Output prior distribution (coloured bars) and kernel density estimates (black line) resulting from samples in subplot A. (C) Posterior distribution over inputs. (D) Output distribution (orange histogram) and target density (blue curve). A Gaussian kernel density estimator with bandwidth 0.05 was fit to the univariate output distribution resulting from 50,000 samples from the bivariate normal prior (using Mathematica’s “SmoothKernelDistribution” function [18]). The posterior distribution was obtained using MCMC implementing the Metropolis algorithm with toroidal boundaries and consisted of 50,000 samples on each of 12 chains. Here 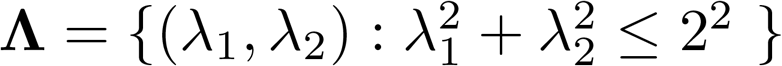.

The posterior distribution that results is located near the bulk of probability mass of the prior distribution (Figure 7C). Comparing this with the distribution that resulted from using a uniform prior (Figure 6C), we see that more informative priors allow us to identify the input parameter distribution. However in both cases the output distribution matched the target (Figure 6D and Figure 7D).

When uniform priors are specified over inputs, the estimated output distribution 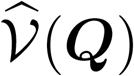 represents estimates of the proportion of total input volume that map to a value of ***Q***. However with non-uniform priors this is no longer true. In this case, we interpret this estimated distribution as representing our *a priori* output distribution that results from our prior weighting over inputs. Here 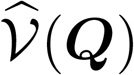 is the estimated probability of obtaining an output ***Q*** given our input priors.

## 5 Results

### 5.1 Inverting classic models of mathematical biology

In order to observe the performance of the CMC algorithm we consider three biological models of increasing complexity.

#### 5.1.1 Logistic growth

We first consider the logistic growth equation,

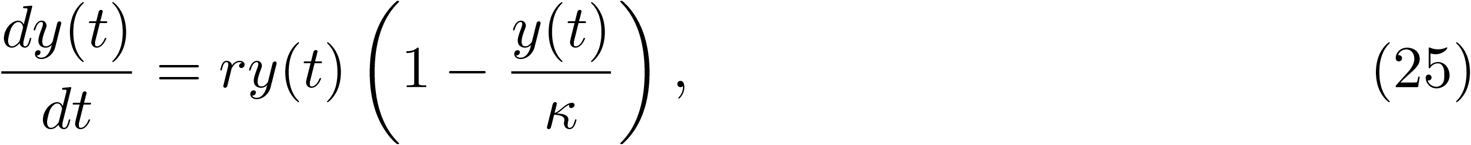

with initial condition

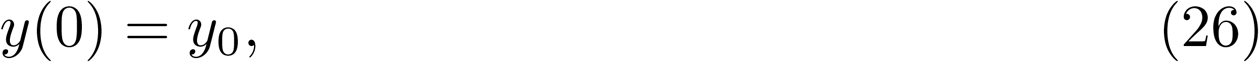

where *y*(*t*) represents the density of individuals at time *t* in a population that experiences competition for resources, for example, a bacterial colony confined to a Petri dish. Here *r* represents the initial (exponential) growth rate of the colony and *κ* is the carrying capacity. An output distribution was generated by calculating *Q*_1_ = *y*(8) for each pair of samples from the following independent input distributions,

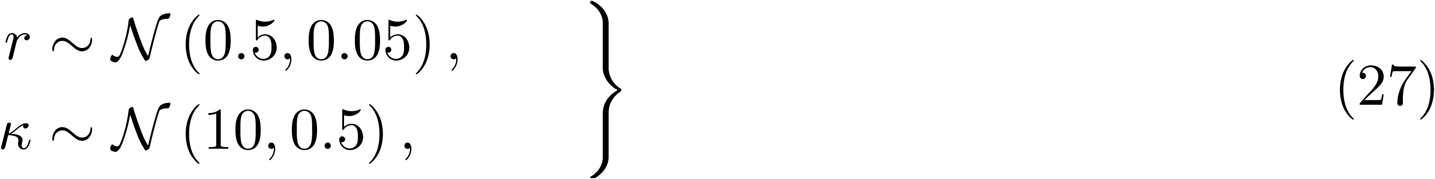

where we assumed that the initial population density was fixed at *y*_0_ = 0.1. The resultant output distribution was fit using a normal distribution and specified as the target. We assumed that the priors for the input parameters were of the form, *r* ~ *U*(0, 2) and *κ* ~ *U*(5, 15), and used CMC to estimate the posterior input distribution. Contour plots of this distribution (Figure 8A) indicate that the growth rate *r* was well identified whereas the carrying capacity was not.

**Figure 8.**
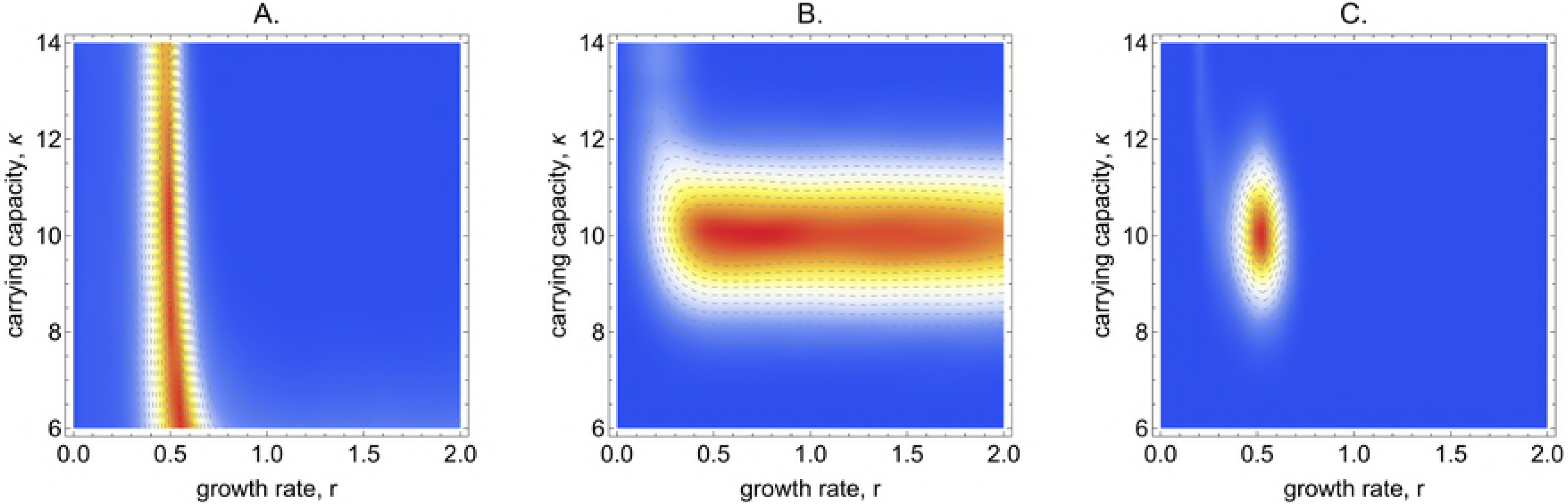
Example §5.1.1: (A) The joint density of growth rate (*r*) and carrying capacity (*κ*) when sampling the population size at *t* = 8. (B) The joint density of growth rate and carrying capacity when sampling the population size at *t* = 29. (C) The joint density of growth rate and carrying capacity when sampling the population size at *t* = 8 and *t* = 29. The contour volumes were estimated using a two-dimensional Gaussian kernel density estimator with default bandwidth fit to 100,000 independent samples from the prior distributions (using Matlab’s “mvksdensity” function [19]), and the posteriors were estimated using 1.2m MCMC iterations (100,000 samples on each of 12 chains).

To explain this pattern of identification, the elasticity of *Q*_1_ with respect to each parameter was calculated using the method described in [20] and is presented in Figure 9. At early times (e.g., *t* = 8), the population density is sensitive to the growth rate parameter *r* but is relatively insensitive to the carrying capacity *κ* since competition for resources has yet to predominate and we are unable to identify this parameter.

**Figure 9.**
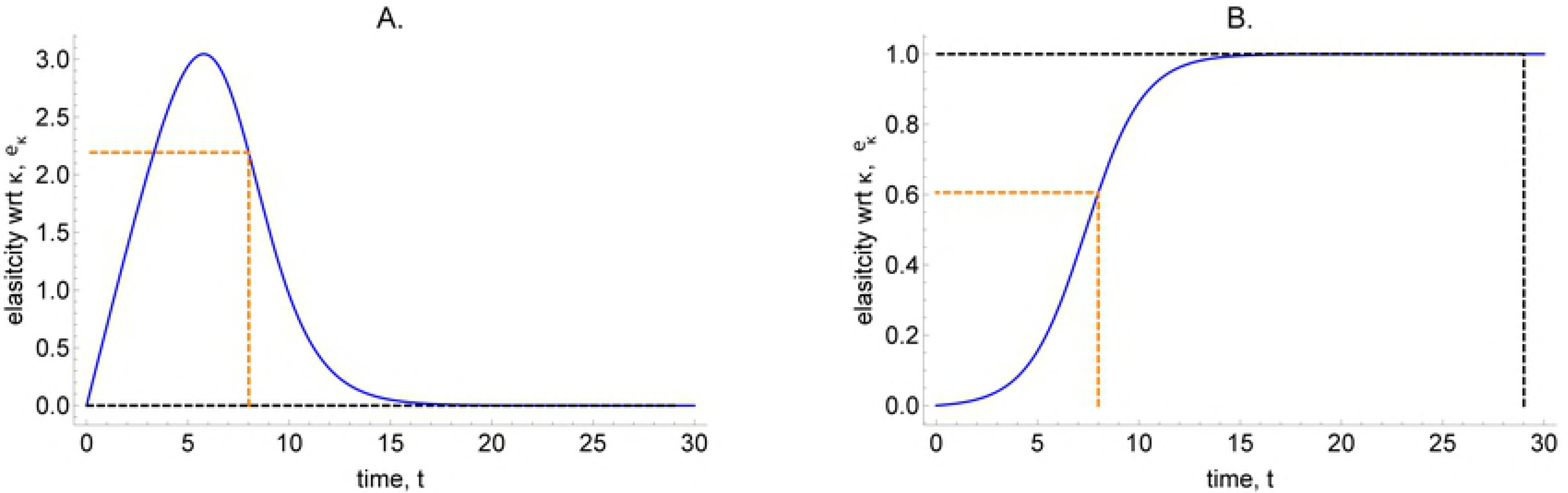
Example §5.1.1: The elasticities of the solution to the logistic equation. (A) Elasticity with respect to growth rate *r*. (B) Elasticity with respect to carrying capacity *κ*. The dashed orange and black lines indicate the elasticities for *Q*_1_ = *y*(8) and *Q*_2_ = *y*(29) respectively. These curves were generated by assuming *y*_0_ = 0.1, *r* = 0.5 and *κ* = 10.

We next chose *Q*_2_ = *y*(29) as the output statistic and used the same input distribution to generate a target output distribution. The posterior input distribution generated by CMC (Figure 8B) identifies the carrying capacity but not the growth rate. At later times (e.g., *t* = 29) the elasticities with respect to the parameters *r* and *κ* are reversed (Figure 9). For *t>* 20, the population density is essentially determined by the (equilibrium) carrying capacity and we are unable to identify the intrinsic growth rate other than to determine it must exceed some minimum value in order for the population to grow to near its equilibrium value by *t* = 29.

Finally we used a target distribution of a bivariate normal distribution of (*Q*_1_*,Q*_2_). The prior distributions were unchanged. The posterior density (Figure 8C) indicates that in this case both input parameters were well-identified. The *combined* distribution of the two previous output statistics is sensitive with respect to each of the parameters. Note that in this case the posterior distribution ascribes probability mass to a region of parameter space that is similar to the original (causative) input distribution. As discussed in §4.1 this is a special case, not typical of underdetermined input-output maps. This is because in the univariate output cases the estimates of the individual parameters are not correlated with one another, and lead to unbiased identification of *r* (for *Q*_1_) and *κ* (for *Q*_2_) respectively. These constraints ensure that using the joint output distribution produces an unbiased estimation of both parameters.

As might be expected, informative priors affect the recovered posterior distribution. To illustrate, we use *Q*_2_ = *y*(29) as our univariate output summary statistic. Given uniform priors, the posterior distribution (Figure 10A and Figure 8B) indicates that, although we can identify the carrying capacity, we cannot identify the growth rate. If instead of a uniform prior, we use *r* ~ Γ(2.5, 0.2) as the prior distribution which allocates probability mass around *r* = 0.5, the resultant posterior narrows with respect to this parameter (Figure 10B). Using a prior distribution that is narrower still, *r* ~ Γ(40, 0.0125), we strongly identify both parameters (Figure 10C).

**Figure 10.**
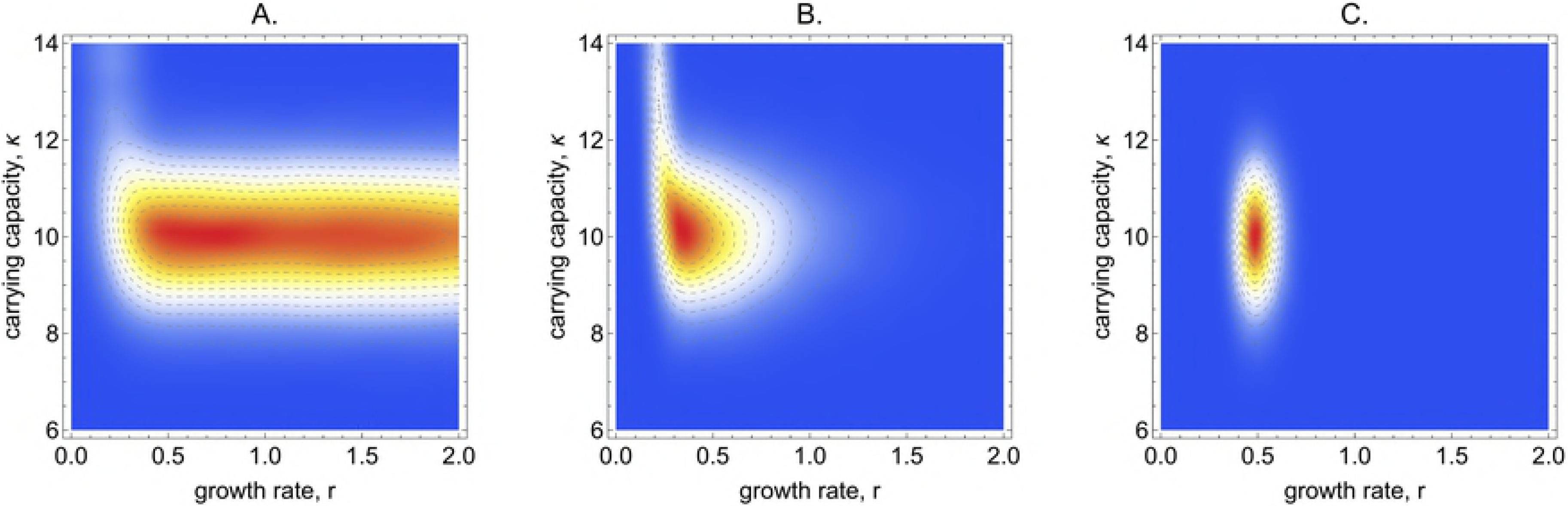
Example §5.1.1: The effect of reducing the variance of the prior distribution for the growth rate on parameter identification. **(A)** *r* ~ *U*(0, 2)**. (B)** *r* ~ Γ(2.5, 0.2)**. (C)** *r* ~ Γ(40, 0.0125). The output distributions were chosen to correspond to a mean of (*r*, *κ*) = (0.5, 10). In all cases the prior specified on the carrying capacity was *κ* ~ *U*(5, 15) and the population size was sampled at time step *t* = 29. The contour volumes were estimated using a two-dimensional Gaussian kernel density estimator with bandwidth 0.05 fit to 100,000 independent samples from the prior distributions (using Matlab’s “mvksdensity” function [19]), and the posteriors were estimated using 1.2m iterations of the Metropolis-Hastings algorithm.

As in Bayesian statistics, there are two ways to identify a parameter: either we collect more data (here by specifying another summary statistic), or we use prior information. In Bayesian statistics, it is the stochasticity of the input-to-output map that generates the uncertainty, meaning that the output distribution (the “data”) can be generated by a collection of possible parameter distributions. In deterministic input-output maps, where the dimension of the inputs exceeds the outputs, it is the multiplicity of the mapping that introduces uncertainty. In both cases, a prior distribution must be specified to ensure that the resultant posterior is unique and a valid probability distribution.

#### 5.1.2 Michaelis-Menten kinetics

The Michaelis-Menten model of enzyme kinetics (see, for example, [8]), describes the dynamics of concentrations of an enzyme (*E*), a substrate (*S*), an enzyme-substrate complex (*ES*), and a product (*P*). Specifically,

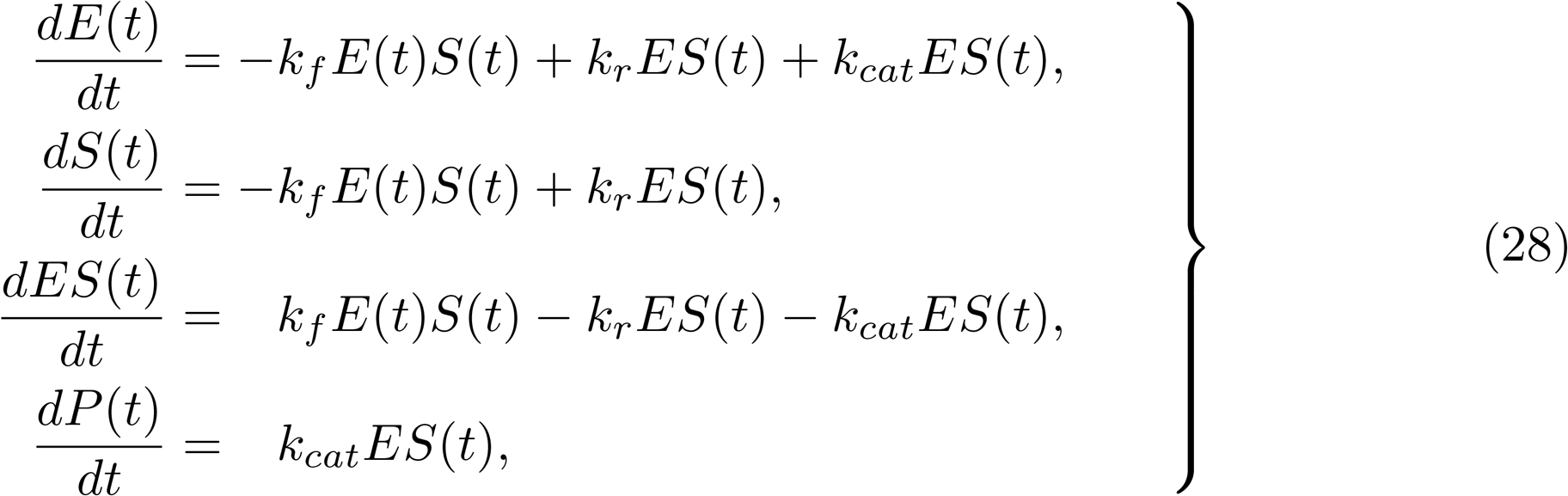

with initial conditions

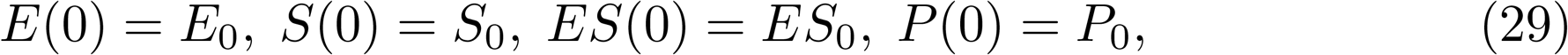

where *k_f_* is the rate constant for the forward reaction *E* + *S* → *ES*, *k_r_* is the rate of the reverse reaction *ES* → *E* + *S*, and *k_cat_* is the catalytic rate at which the product is formed by the reaction *ES* → *E* + *P*. We assume independent uniform priors on each of the parameters (*k_f_,k_r_,k_cat_*), with upper and lower bounds indicated by the axes in Figure 11. We target the bivariate output distribution given by,

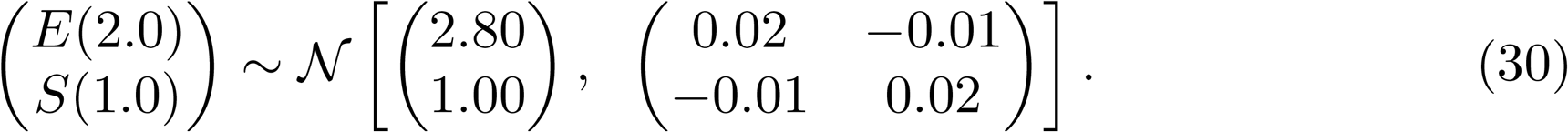

**Figure 11.**
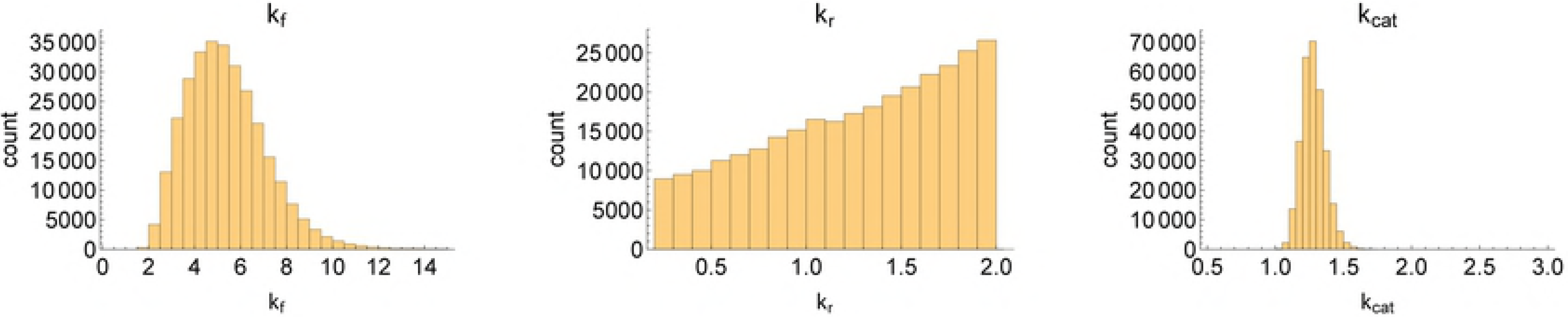
Example §5.1.2: The marginal densities of parameters for the Michaelis-Menten model. The parameters were assigned uniform priors over the ranges shown in the axes. The contour volumes were estimated using a Gaussian kernel density estimator with default bandwidth fit to 120,000 iterations (using the “kde” function in the “ks” R [21] package with grid sizes of 300 and maxima of 4 in each dimension [22]), and the posteriors were estimated using c.300,000 MCMC iterations.

Running CMC resulted in output distributions which matched the target (see S1–Michaelis Menten). The posterior input distribution indicates that *k_cat_* is strongly identified by this choice of output distribution and *k_f_* is restricted to a subset of its prior bounds (Figures 11 & 12). The rate of the backward reaction *k_r_*, however, cannot be recovered from this output target.

**Figure 12.**
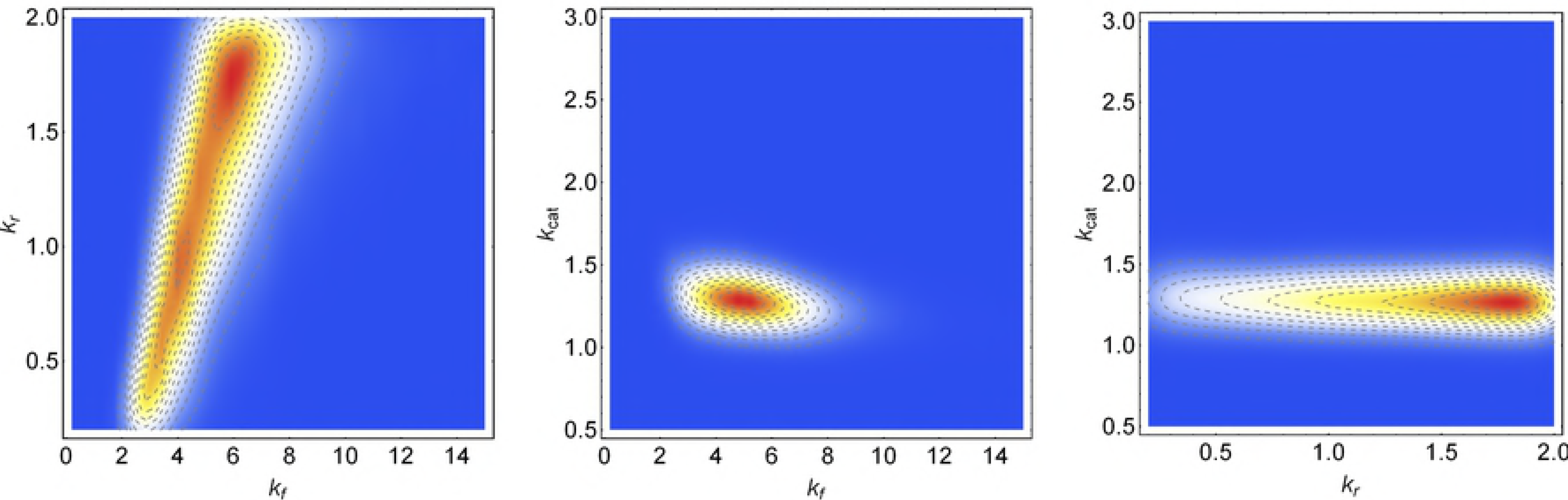
Example §5.1.2: The joint distribution of pairs of parameters for the Michaelis-Menten model. The parameters were assigned uniform priors over the ranges shown in the axes. The contour volumes were estimated using a Gaussian kernel density estimator with default bandwidth fit to 120,000 iterations (using the “kde” function in the “ks” R [21] package with grid sizes of 300 and maxima of 4 in each dimension [22]), and the posteriors were estimated using c.300,000 MCMC iterations.

Some insight in to this pattern of identifiability can be gained by examining the elasticities of *E an*d *S t*o each of the input parameters (Figure 13). The enzyme and substrate concentrations at times *t* = 2 and *t* = 1, respectively, are most sensitive to changes in *k_cat_*, and this parameter is the most clearly identified. The enzyme and substrate concentrations at our sampling times are less sensitive to the forward rate of reaction, k*_f_*, but these values are apparently sufficient to localise this parameter to a subregion of its prior bounds (Figure 11). The rate of the backward reaction, however, does not strongly influence either the sampled values of the enzyme or substrate concentrations at our sampling times, and this parameter is practically unidentifiable.

**Figure 13.**
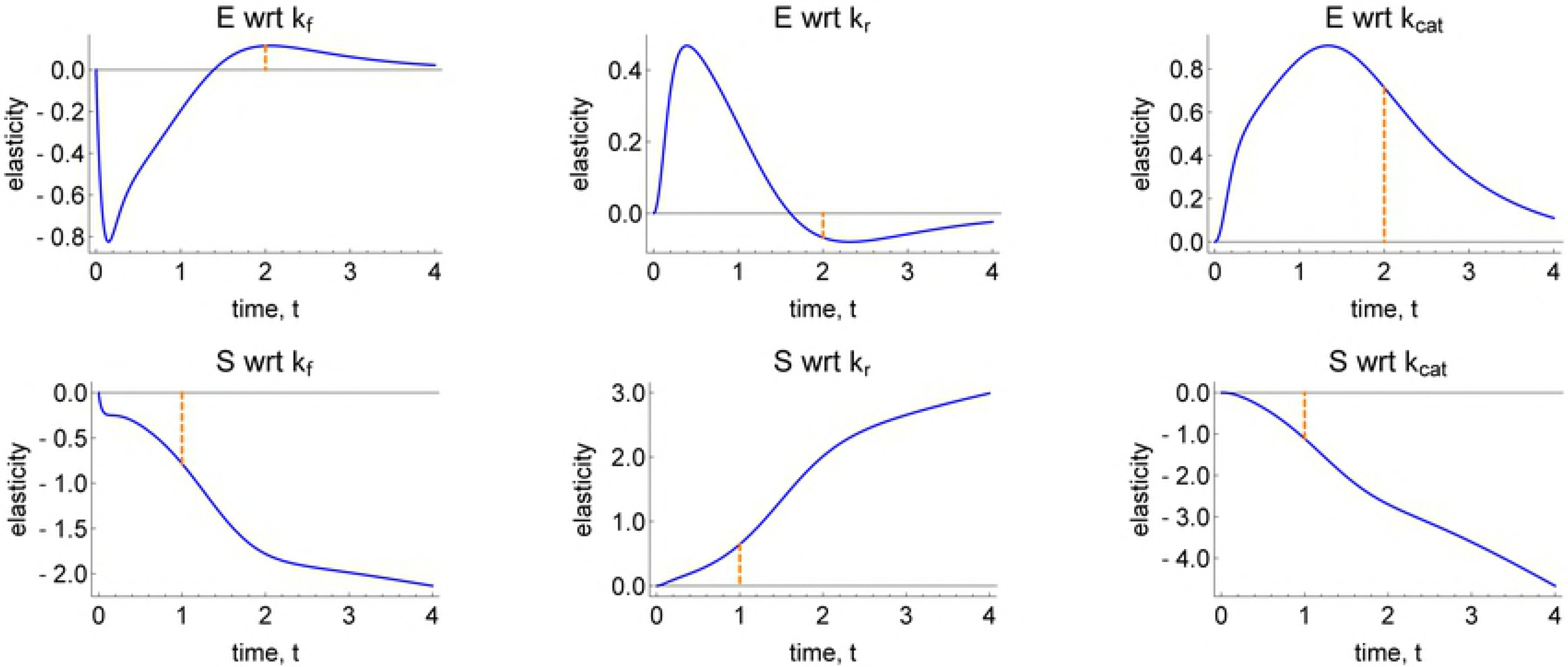
Example §5.1.2: The elasticities of the enzyme (top row) and substrate (bottom row) concentrations with respect to the three input parameters. The dashed orange lines indicate the times at which each output is sampled in our example. These curves were generated by assuming *k_f_* = 2, *k_r_* = 1, *k_cat_* = 1.5, *E*(0) = 4, *S*(0) = 8, *ES*(0) = 0, and *P* (0) = 0.

#### 5.1.3 SIR model

An SIR model of disease transmission incorporating a carrying capacity for the population is described by the following system of ODEs [23],

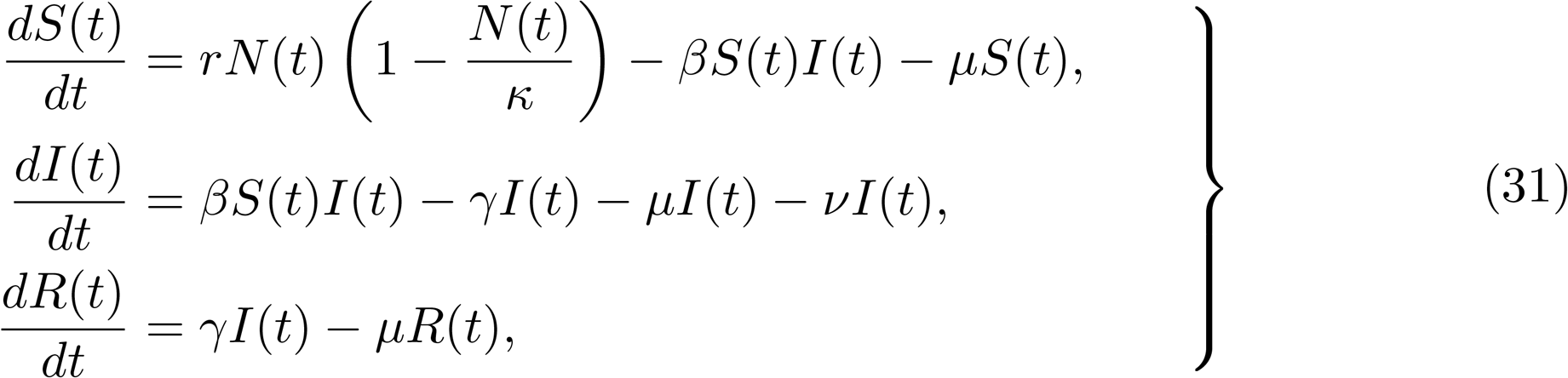

with initial conditions

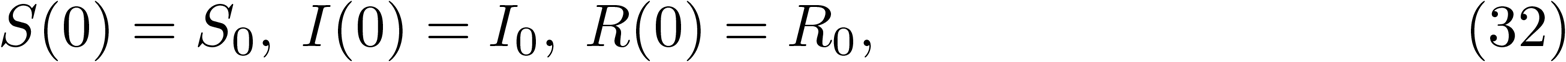

where *N*(*t*) = *S*(*t*)+ *I*(*t*)+ *R*(*t*) is the total population density at time *t*.

Including the initial conditions (*S*_0_*,I*_0_*,R*_0_), we have nine uncertain parameters, each of which is assumed to have uniform prior distributions. The upper and lower bounds of these distributions are provided by the axes in Figure 14. We use a three dimensional output statistic in which the elements consist of sampling the infected population at *t* = 10 and the recovered population at times *t* = 10 and *t* = 80. For our target distribution, we choose the following multivariate normal,

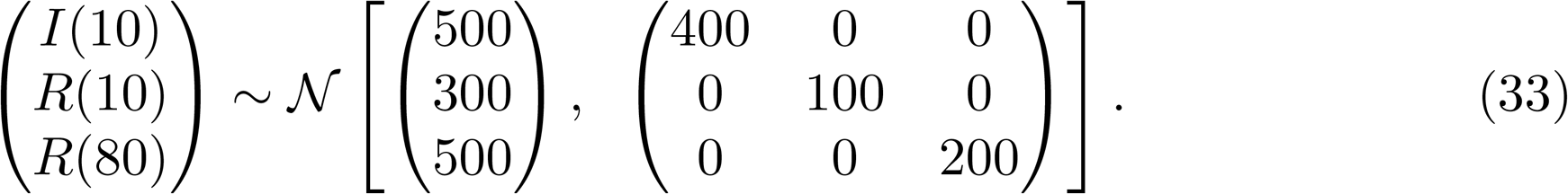

**Figure 14.**
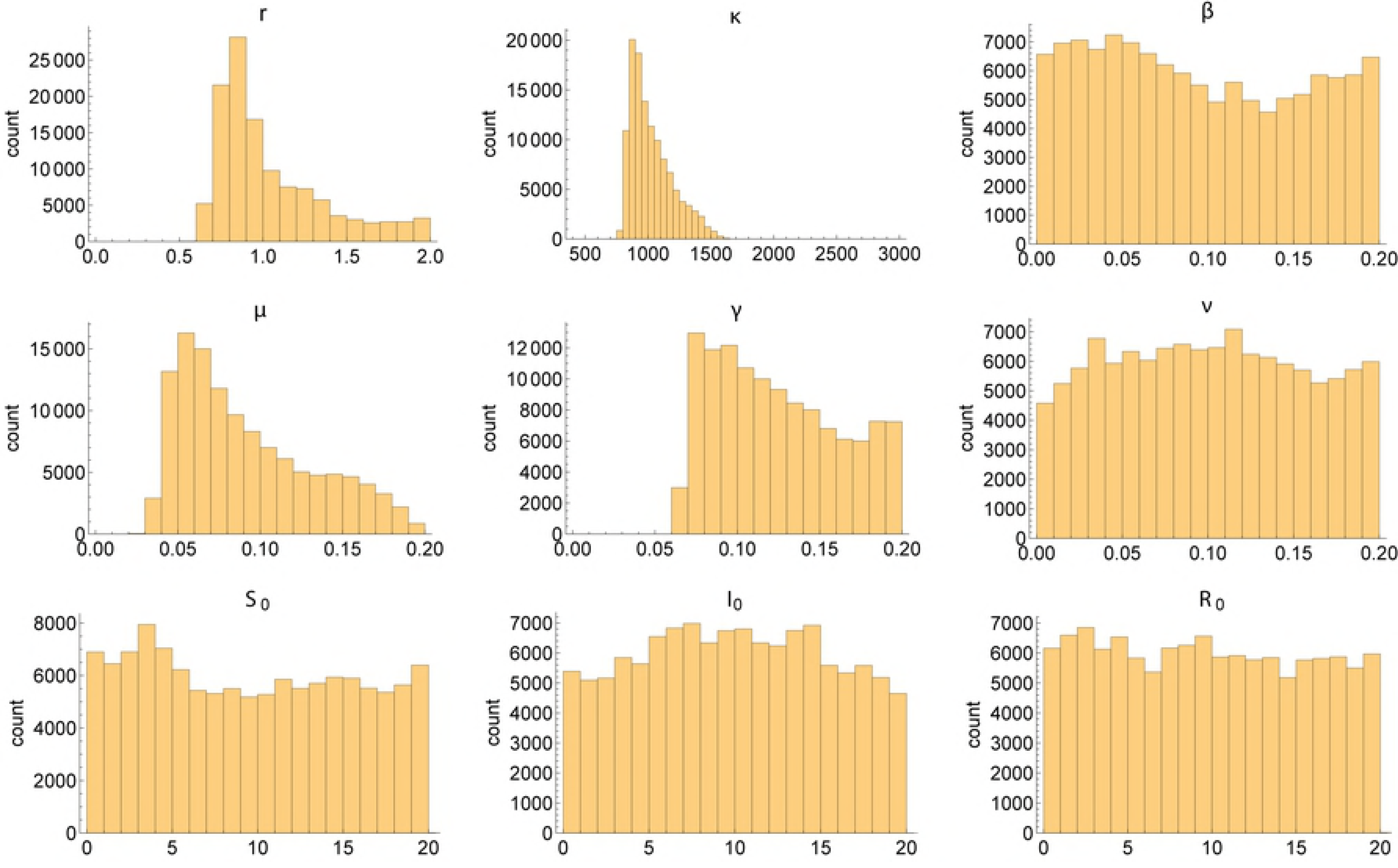
Example §5.1.3: The marginal densities of parameters for the SIR model. The parameters were assigned uniform priors over the ranges shown in the axes. The contour volumes were estimated using a Gaussian kernel density estimator with default bandwidth fit to 100,000 iterations (using the “kde” function in the “ks” R [21] package with grid sizes of 300 and maxima of 3000 in each dimension [22]), and the posteriors were estimated using 1.2m MCMC iterations (100,000 samples on each of 12 chains).

We restrict the dimension of the output space in order to examine a highly underdetermined system and to avoid issues associated with kernel density estimation in higher dimensional output spaces. A further discussion of this issue appears in §6.

The posterior distribution over inputs indicates modest identification of the parameter *r*, which dictates the rate of initial exponential growth for the susceptible population (Figure 14*r*) and the carrying capacity term *κ*. The posterior distribution of the death rate *µ* and of the recovery rate *γ* was also constrained to a subset of input space (Figure 14*κ*, 14*µ*, 14*γ*). The remaining parameters of the model were unidentified using these output statistics (Figures 14). For examples of joint distributions of selected pairs of parameters see S2–SIR model. For the posterior input distribution that resulted, we calculated the corresponding output distribution, and compared this with the marginal target distributions, indicating good agreement (see S2–SIR model).

## 6 Discussion

Natural selection has dictated that organisms have often evolved redundancies for systems essential for life. By building mathematical models we aim to mimic these systems, meaning that these models should embody similar redundancies as their biological counterparts. Assessing the sensitivity of key characteristics of model outputs to perturbations in input parameters provides insight into the sensitivity of the system to each of its constituent elements. These so-called sensitivity analyses of mathematical models allow us to probe the biological system even when biological experiments are infeasible. Inverse sensitivity analysis inverts this process and instead of determining how model outputs vary in response to changes to the input parameter values, estimates a distribution over inputs which achieves a given distribution over outputs.

In this paper, we introduce an approach to inverse sensitivity analysis which can be applied to systems with many input parameters, mitigating the curse of dimensionality that limits the scope of methods which rely on grid-based approaches to build an explicit output-to-input map [10]. We have demonstrated that our algorithms can perform inverse sensitivity analysis on mathematical models of biological systems across a range of complexities, including the logistic growth model (2 inputs & 1 output target), Michaelis-Menten kinetics (3 inputs & 2 output targets), and the SIR model with uncertain initial population sizes (9 inputs & 3 output targets). As well as detailing our algorithms, we provide a probabilistic framework for understanding inverse sensitivity analysis, in which “prior” probability distributions are set on the inputs. These prior beliefs over input values are consistent with a “posterior” input distribution which, when transformed through the input-to-output map, results in a “target” output distribution. To sample from these posterior input distributions, we introduce a two-step sampling algorithm. In the first of these steps, input parameters are independently sampled from their prior distributions and, by fitting a kernel density estimator to the output values, this provides an approximate Jacobian transform (which we interchangeably term a “contour volume distribution”), which is used in the second step involving Markov chain Monte Carlo. A similar algorithm for inverse sensitivity analysis has recently been derived from a measure theoretic perspective by [24]. These authors also investigate stability of the posterior distribution with respect to the observed output distribution, the assumed prior distribution and the approximation of the contours of the forward map. We believe that the different path we take to the shared goal offers complementary insight into the algorithm’s mechanism and provides an intuitive way to understand inverse sensitivity analysis, more generally.

There are several subtleties in the first steps of the processes described in Algorithms 1 and 2) which must be understood in order to ensure a valid input distribution is obtained. Indeed, these intricacies complicated our own efforts in testing the algorithms. Provided the output is well-behaved over the space of possible input values, a univariate output distribution can be approximated given a relatively modest number of samples from the input priors using standard kernel density estimation (KDE). Here, for the univariate output target distributions we found that KDE with a Gaussian kernel using default bandwidths from each software package used (Matlab, Mathematica and R [18, 19, 21]) was able to represent the output distribution with sufficient fidelity to ensure the input posterior recaptured the output target. The number of input samples necessary to ensure convergence to the true posterior input distribution, however, depends on the exact output distribution being targeted. If the bulk of probability mass for the target output distribution lies at a location in output space where the contour volume is rapidly varying, then the input distribution obtained will be sensitive to errors in kernel density estimates of the contour volume distribution, and many samples will be required. Similarly, if a region of low contour volume is targeted, then kernel density estimates with few samples will be relatively noisy and more samples will be necessary. Here we have assumed numerical errors in solving the map are negligible and independent of the parameters. Neither assumption is likely to be true for sophisticated partial differential equation models and the interaction between numerical and sampling errors is the subject of ongoing analysis. KDE introduces a further source of error, which must be carefully managed to ensure reasonable results are obtained.

Our algorithms avoid the curse of dimensionality of the input space which plagues grid-based approaches to inverse sensitivity analysis. The necessity of having to fit a probability distribution to the output samples resultant from sampling the prior input distributions means that, at present, our approach is limited to problems with relatively few outputs. In §5.1.3, the output target was a three-dimensional distribution, and we expended considerable effort finding a KDE method that adequately approximated the three-dimensional contour volume distribution. We ultimately found that the most effective approach was obtained by using the “kde” function within the “ks” R package [21, 22], which uses the data to estimate unconstrained bandwidth matrices, which are then used to fit kernel density estimates to data with up to six dimensions. Density estimation, however, is currently an active area of research and software packages exist implementing many different variants of KDE (see [25] for a review of the R packages already available in 2011). Vine copulas have recently been suggested as an approach which avoids the curse of dimensionality in density estimation [26]. If this promise is realised, then our algorithm will be applicable to output target distributions of higher dimensions.

Mathematical models have proved indispensable tools for elucidating understanding of biological systems, which are frequently not amenable to direct experimentation. Biological systems are often robust to perturbations to particular constituent processes, and we can use mathematical models to explore these sensitivities. Inverse sensitivity analyses are a relatively recent addition to a modeller’s toolbox, which allows one to determine an input distribution - consistent with prior beliefs - that can generate a given distribution of outputs. Here we introduce a Monte Carlo method which extends the range of models for which inverse sensitivity analysis can be performed, and illustrate its utility for several problems of interest to computational biology. It is our hope that, by publishing this method, others are encouraged to undertake inverse sensitivity analysis, which we have found is insightful for building and analysing mathematical models.

## 7 Author contributions

BL, DJG and SJT conceived the study. BL and SJT carried out the analysis. All authors helped to write and edit the manuscript.

## Supporting information captions

- S1–Michaelis-Menten kinetics
- S2–SIR model
- S3–A nonlinear map

